# Phenotype-first covalent fragment screening identifies a synthetic lethal TYMS inhibitor in ATRX-deficient cells

**DOI:** 10.64898/2026.06.10.731339

**Authors:** Federica Raguseo, Elliot Fellows, Cesira de Chiara, Benjamin J. Mortishire-Smith, Sandra Segura-Bayona, Emma Cawood, Ming Jiang, Fernanda Teixeira Subtil, Aurora I. Idilli, William J. McCarthy, Michael Howell, Katrin Rittinger, James I. MacRae, Mark Skehel, Andrew J. Powell, David House, Jacob T. Bush, Simon J. Boulton

## Abstract

Chemoproteomic mapping of covalent fragment libraries is expanding the ligandable human proteome with direct evidence of cellular target engagement. However, understanding the functional consequences of specific covalent modifications typically requires extensive downstream biological characterisation. Here we present a ‘phenotype-first’ approach that integrates covalent fragment screening with chemoproteomics and genetic deconvolution in a disease-relevant context. Using isogenic ATRX wild-type and knockout eHAP iCAS9 cells, we screened a library of around 500 cysteine-reactive fragments for differential cell killing and identified a chloroacetamide fragment, PP12, that selectively impairs the viability of ATRX-deficient cells. By combining competitive click-chemoproteomics with genome-wide CRISPR synthetic lethal datasets, we identified thymidylate synthase (TYMS) as a phenotypically relevant target of PP12. Target validation was supported by crystallography, competition with the active-site inhibitor 5-fluorouracil, and impaired dTMP synthesis in cells. Mechanistically, TYMS inhibition induces replication stress that is selectively cytotoxic to ATRX-deficient cells and is dependent on FAM111A and SLFN11. This work establishes a generalisable workflow linking covalent fragment phenotypes to target deconvolution using chemoproteomics and mechanistic validation.

## INTRODUCTION

Over the past decade, CRISPR–Cas (Clustered Regularly Interspaced Short Palindromic Repeats–CRISPR Associated Protein) screening has enabled the systematic identification of genes linked to specific phenotypes across diverse, disease-relevant models^1^. In cancer biology, these approaches have been widely applied to uncover biomarkers and genetic interactions that can inform new therapeutic strategies^2,3^. Genome-wide CRISPR screens have been particularly powerful for probing synthetic lethality (SL), in which selective cell death is achieved by exploiting vulnerabilities that emerge in cancer cells^4^. In a SL interaction, a target gene is essential in cancer cells (where partner genes may be mutated or downregulated) but is dispensable in normal cells^5,6^ (**Fig. 1a**). Clinically relevant synthetic lethal pairs therefore offer opportunities to selectively target cancer cells while minimizing off-target toxicity in normal tissues^5,7^. For example, the chromatin regulator ATRX has been reported to exhibit SL interactions with multiple genes across several studies, including genome-wide CRISPR knockout screens^8,9^ and focused drug screens (*e.g.* ATRX–PARP SL)^10^. Given the large number of reported ATRX SL partners^8^ and the fact that ATRX is among the most frequently mutated proteins across cancers^11,12^, ATRX deficiency represents a promising context for therapeutic target discovery^13^. However, translating genetic vulnerabilities into therapeutics remains a bottleneck, as the identification of tractable binding sites on SL partners and development of suitable small-molecule starting points is often challenging and resource intensive^14^.

**Fig. 1.**
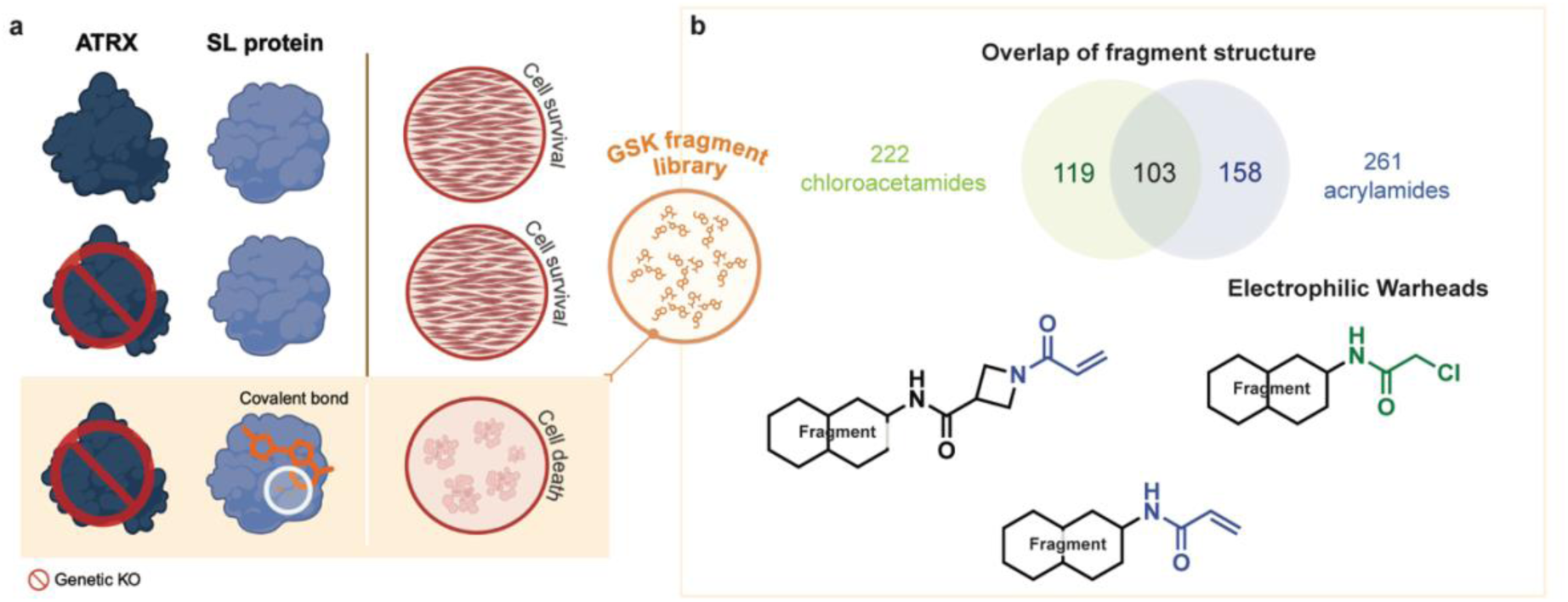
Phenotype-First reactive fragments screening platform. **A.** *Schematic for the Synthetic Lethality (SL) phenotype.* ATRX-deficient cells are viable unless a SL pathway is perturbed by a covalent fragment. **B**. *Overview of the reactive fragment library screened.* The library comprises 222 chloroacetamides and 261 acrylamides; some fragment structures are common to both electrophiles.

Covalent fragment–chemoproteomics screening has emerged as a useful strategy for identifying small molecule binders across the proteome, expanding the set of proteins for which actionable chemical starting points are available, including targets historically considered difficult to drug^15,16,17^. This strategy offers two complementary advantages. First, covalent libraries can enable potent, durable target engagement in cells across diverse protein classes, even with modest library sizes, by leveraging the efficient chemical-space sampling of fragments^18^. Second, chemoproteomics provides a direct and sensitive readout of target engagement in biologically relevant live-cell settings, enabling systematic mapping of ligandable sites in complex proteomes^19^. Cysteines residues are the most commonly targeted residues in covalent ligand discovery, reflecting both their high nucleophilicity and the extensive precedent of cysteine-targeting covalent drugs and chemical probes^20^. In a landmark study, Backus *et al.* screened a covalent fragment library in cell lysates and identified ligands for more than 700 cysteines in the human proteome^16^, including sites on proteins previously considered difficult to drug^21^. Subsequent screens in both cells and lysates have broadened the druggable proteome by pairing complementary methods to directly assess target engagement, including mass spectrometry (MS) and tailored functional assays^22,23^.

Identifying a ligandable cysteine, however, does not guarantee functional consequence. A key limitation of covalent fragment screening by chemoproteomics (‘chemoproteomics-first’) is that demonstrated target engagement at a specific residue does not necessarily translate into functional modulation or an observable phenotype^23^. Although covalent binding can compensate for the low affinity inherent to fragment interactions, even potent and selective covalent ligands may be without functional consequence if they bind outside regions critical to protein function. A direct route to circumventing this limitation is to screen fragments directly in cells for phenotypes of interest using a ‘phenotype-first’ approach that is inherently target-agnostic. In this framework, hits are prioritised based on cellular activity, and subsequent chemoproteomics target deconvolution can reveal therapeutically relevant targets and unanticipated biological mechanisms^24,25^.

Here, we integrate ‘phenotype-first’ covalent fragment screening with chemoproteomics-based target deconvolution and insights from genome-wide CRIPSR-Cas9 drop-out screening. Using ATRX deficient isogenic systems, we identified a covalent fragment with SL activity and linked this phenotype to its molecular target and mechanism of action.

## RESULTS

### Identification of PP12, a covalent fragment inducing synthetic lethality in ATRX^KO^ cells

Human chronic myeloid leukaemia (eHAP) cell lines expressing either wild-type (WT) or ATRX knockout (ATRX^KO^) inducible Cas9 (iCAS9) were screened in parallel against a cysteine-reactive fragment library comprising 222 chloroacetamides and 261 acrylamides (**Fig. 1b, Fig. S2**). Compounds within this library were selected in a way that maximised chemical diversity, aligned to traditional fragment ‘rule of three’ parameters: <300 Da molecular weight and ≤3 hydrogen-bond donors and acceptors. We hypothesized that fragments engaging proteins synthetically lethal with ATRX would selectively reduce viability of ATRX^KO^ cells relative to WT cells (**Fig. 1a**). Compounds were tested at three concentrations (50, 10 and 1 µM) over a 4-day exposure window, and cell viability was quantified by nuclei count (**Fig. 2a**).

**Fig. 2.**
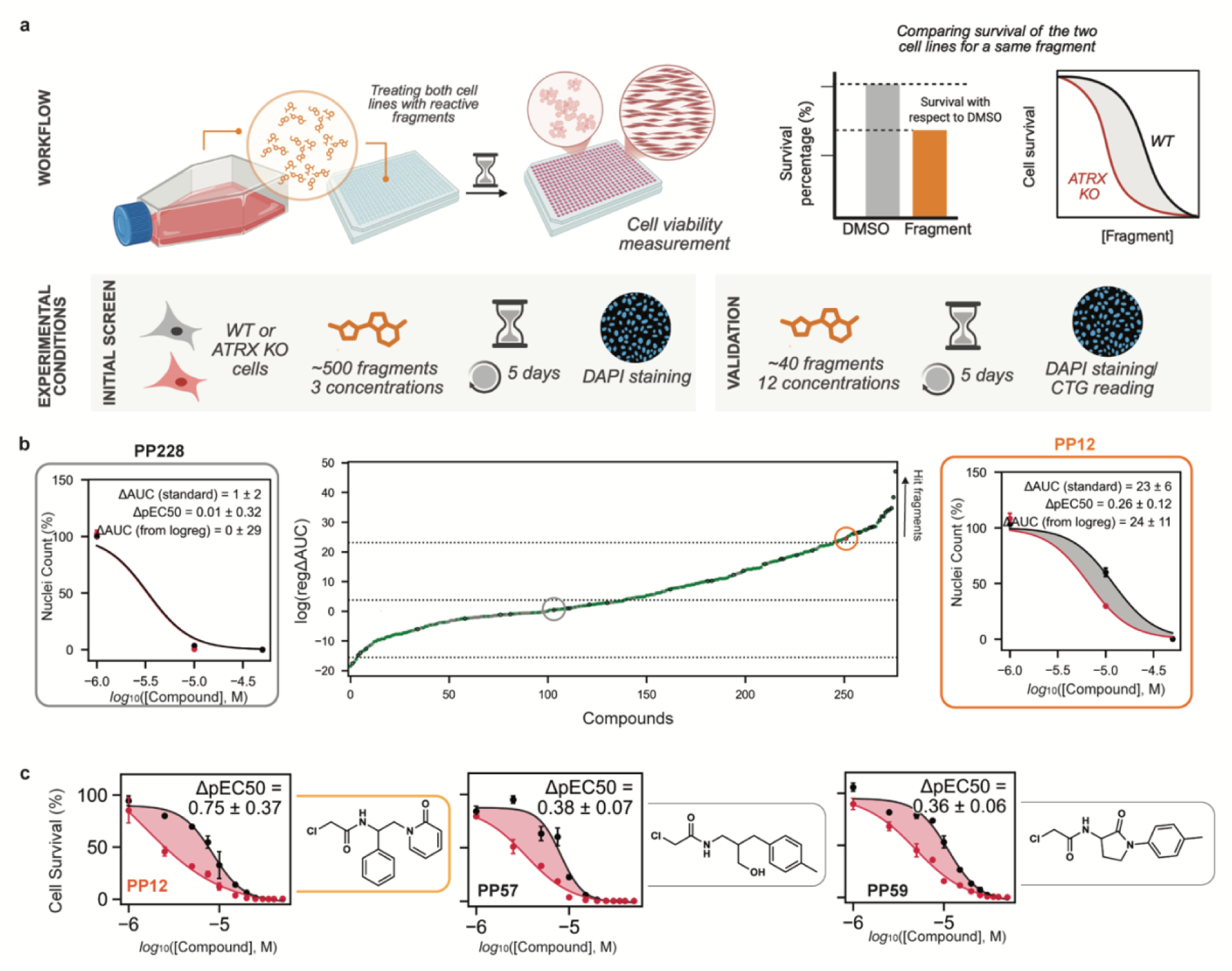
High-Throughput Synthetic Lethality Screen for fragment ID. **A.** Overview of screen and confirmation. WT and ATRX^KO^ cells were treated with 3 fragment concentrations for 4 days and viability was quantified via DAPI nuclear staining. The phenotype was quantified by normalizing to DMSO controls and comparing survival between WT and ATRX^KO^ cells. Confirmation used the same experimental workflow with expanded concentration range. **B.** Fragments ranking by SL phenotype (ΔAUC). Points show ΔAUC ranking for chloroacetamides (green) and acrylamides (grey) passing filtering criteria (**Fig. S1a**). Fragments for concentration-response confirmation are outlined. PP12 is highlighted. Representative curves are shown for inactive fragment PP228 (LHS) and PP12 hit compound (RHS) N=3. **C.** Confirmation curves and structures for the top three hit fragments N=3.

For each fragment, three-point concentration–response data were fitted with a restrained logistic model to estimate area under the curve (AUC) (**Fig. 2b**, **Fig. S1a**). Fragments were then ranked by differential viability between WT and ATRX^KO^ cells (ΔAUC) (**Fig. 2b**, **Fig. S1a**). Larger ΔAUC values indicated greater selective toxicity toward ATRX^KO^ cells, whereas smaller ΔAUC values reflected little to no differential effect (**Fig. 2b**). We defined hits using an absolute ΔAUC threshold of ≥ mean ± 2×SD (threshold = 23), yielding 28 hit fragments (∼6% hit rate). Compounds of interest were validated in 12-point concentration–response experiments, along with structurally similar equivalents and non-hits negative controls (**Fig. S1b**).

The SL phenotype in 12-point concentration response was quantified using ΔpEC_50_ (WT - ATRX^KO^), where pEC_50_ refers to the log_10_ concentration at which viability was reduced by 50%. The three fragments that gave the highest ΔpEC_50_ were prioritised for follow-up (PP57, PP59 and PP12), with PP12 emerging as the highest-ranking fragment with a ΔpEC_50_ of 0.75 ± 0.37 (**Fig. 2c**).

### PP12 activity is reproducible across clones and cell lines and supports functionalisation for chemoproteomics

To confirm the SL phenotype observed with PP12, we measured viability by CellTiter-Glo assay, in a second ATRX^KO^ clone generated with a distinct single guide RNA (sgRNA) (g4) to mitigate potential clonal effects^26^. We also tested PP12 in an additional isogenic pair (human lung cancer NCI-H460 iCAS9 WT/ATRX^KO^) to evaluate activity across a distinct genetic and molecular background^27^ (**Fig. 3a, 3b**). In both settings, PP12 activity was benchmarked against AZD6738^28^, an ATR inhibitor (ATRi, Ceralasertib) previously reported to be synthetic lethal with ATRX deficiency^8^. To determine whether cellular activity depended on covalent engagement, we also tested a ‘non-covalent’ analogue of PP12 (PP2206) where the chloroacetamide moiety was replaced with an isosteric propanamide. PP2206 showed no phenotypic effect (**Fig. 3c**), indicating that covalent reactivity via the chloroacetamide moiety is required for target engagement in cells.

**Fig. 3.**
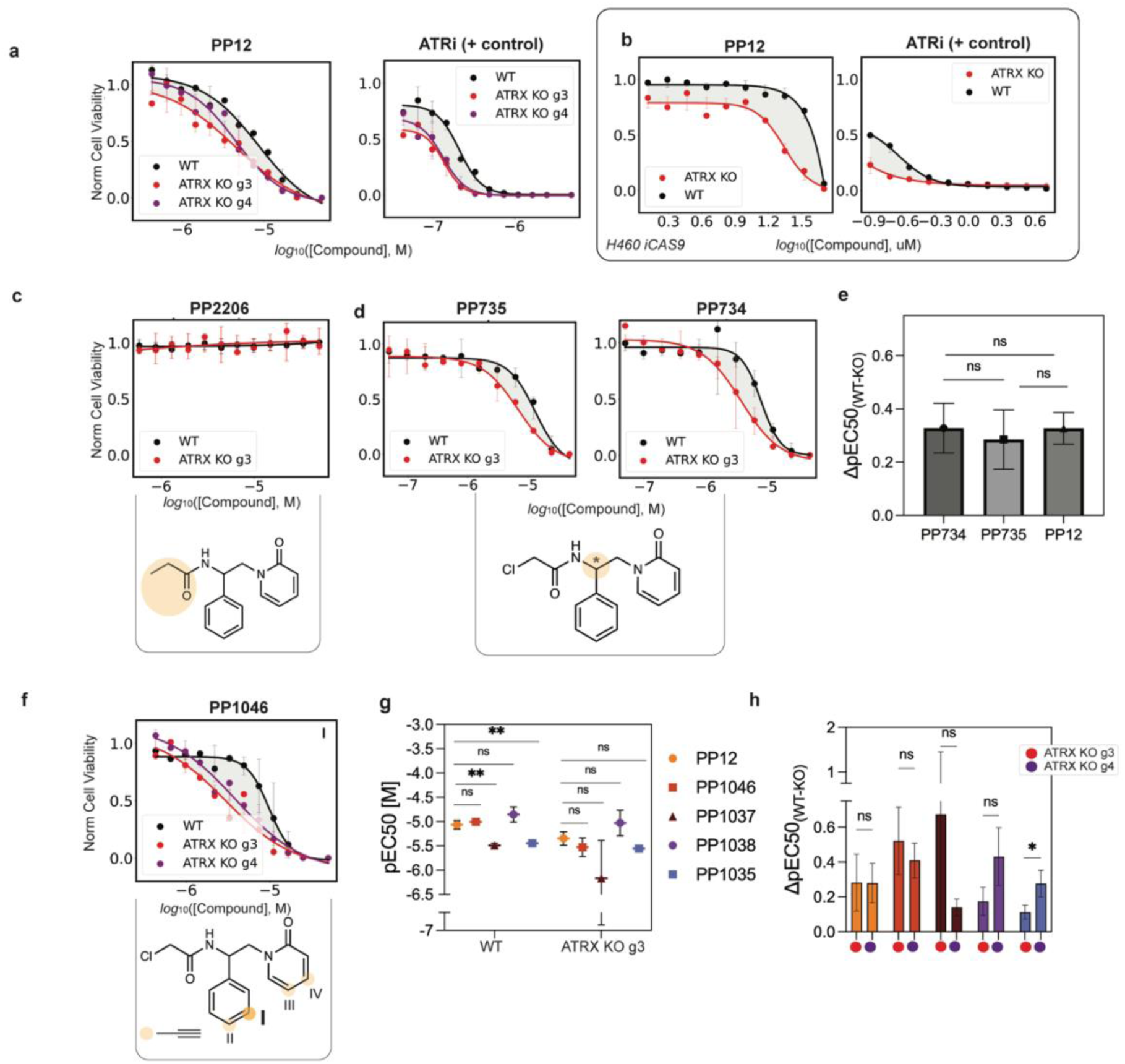
Hit Fragment Validation and selection of a clickable analogue. **A.** PP12 concentration-response in eHAP ATRX^KO^ g3 and g4 (CTG) N=3. Curves for other hit fragments shown in **Fig. S2**. **B.** PP12 concentration-response in H460 ATRX^KO^ (CTG) N=3. **C.** PP2206 (non-covalent control) shows no cell viability effect (CTG) N=3. **D.** Stereoprobes PP735 and PP734 concentration-response in eHAP ATRX^KO^ (CTG) N=3. **E.** No statistically significant differences observed between PP12, PP735 and PP734 (*P*-values (PP734-PP735: 0.64, PP12-PP734: 0.99, PP12-PP735: 0.60) - calculated by unpaired t-test) N=3. **F.** PP1046 (click-derivative) concentration-response in eHAP ATRX^KO^ (CTG in both ATRX^KO^ g3 and g4) N=3. **G.** pEC_50_ comparison of PP12 and click-derivatives (*P*-values in WT PP1046: 0.34, PP1037: 0.002, PP1038: 0.11, PP1035: 0.002). *P*-values in ATRX^KO^ (PP1046: 0.25, PP1037: 0.15, PP1038: 0.14, PP1035: 0.06 calculated by unpaired t-test) N=3. **H.** Click-derivatives ΔpEC_50_ (WT-KO) compared between both ATRX^KO^ clones. *P*-values (PP12: 0.98, PP1046: 0.42, PP1037: 0.30, PP1038: 0.07, PP1035: 0.03) – calculated by unpaired t-test) N=3.

Stereoprobes have often been used to identify ligand–protein interactions that are sensitive to absolute stereochemistry^29,30^. We therefore separated racemic PP12 by chiral chromatography into its two enantiomers (PP734 and PP735). Both enantiomers were tested in eHAP WT and eHAP ATRX^KO^ cells with no statistically significant difference in the SL phenotype as quantified by ΔpEC_50_ (WT-ATRX^KO^) (**Fig. 3d, e**), indicating that engagement of the phenotype-driving target is not enantioselective in this case.

To enable target identification by click-chemoproteomics, we designed a set of alkyne-functionalised PP12 analogues. Four racemic analogues (PP1046, PP1037, PP1038 and PP1035) were evaluated in concentration–response format over 5 days in WT and ATRX^KO^ cells (**Fig. 3f**, **Fig. S3a**). Analogues were compared with PP12 based on (i) similar potency in WT and ATRX^KO^ eHAP cells (**Fig. 3g**), and (ii) clone-independent SL across both ATRX^KO^ clones (**Fig. 3h**). Although all analogues produced some degree of synthetic lethality, PP1046 best satisfied these criteria and was selected for subsequent chemoproteomics experiments.

### Chemoproteomics identifies TYMS as a target of PP12

To identify the protein targets of PP12, we performed competitive click-chemoproteomics in live cells. Briefly, eHAP ATRX^KO^ cells were pretreated with DMSO vehicle or increasing concentrations of PP12 (competitor) to block covalent binding sites prior to addition of the alkyne probe PP1046. Following CuAAC (Copper-catalysed Azide–Alkyne Cycloaddition) with biotin-azide and subsequent affinity enrichment, PP1046-bound proteins were digested and peptides analysed by LC–MS/MS (**Fig. 4a**, **Fig. S3a**)^31–33^. Proteins showing concentration-dependent competition by PP12 were considered as candidate targets for the SL phenotype.

**Fig. 4.**
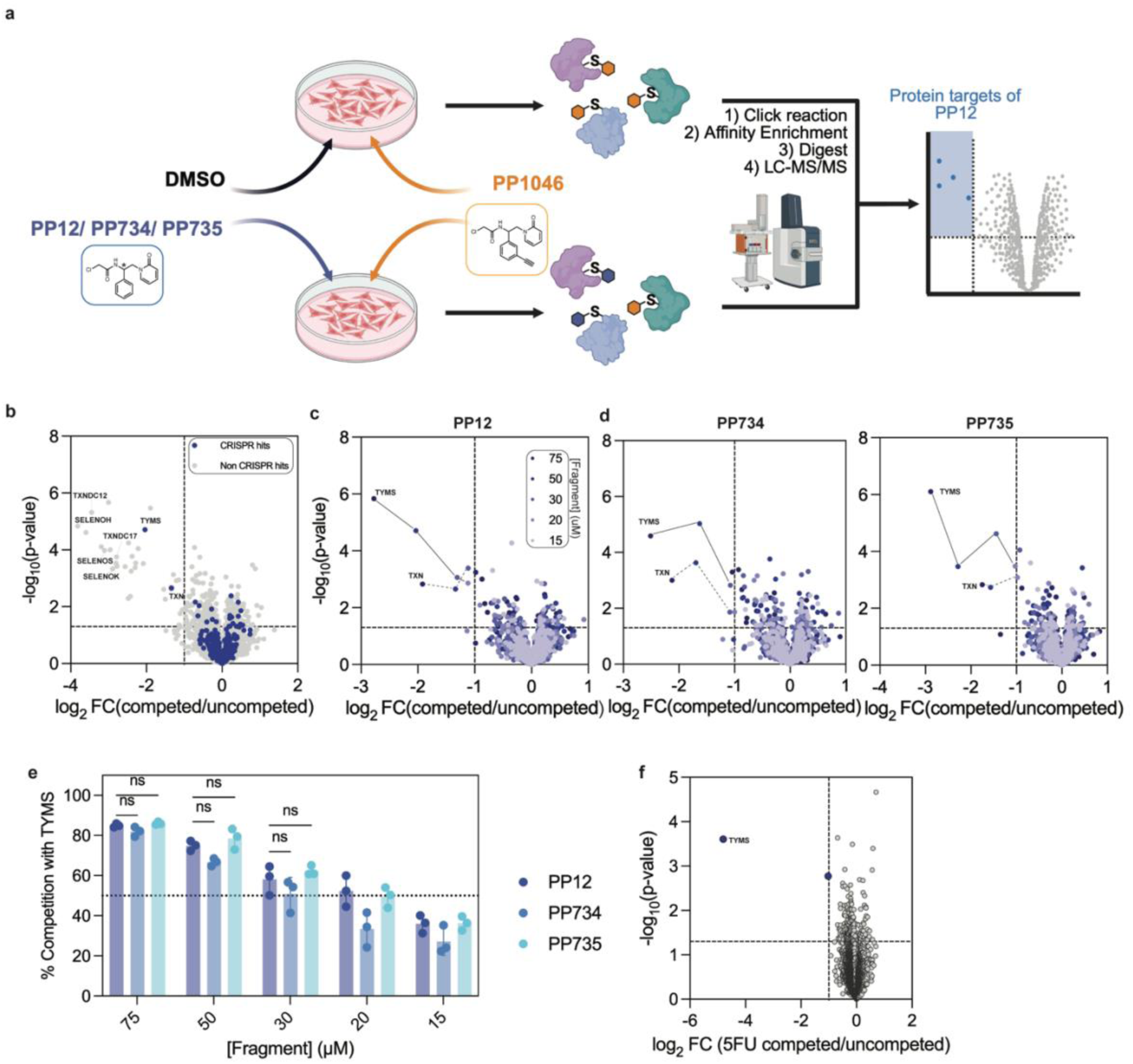
TYMS target ID via chemoproteomics approaches. **A.** Competitive click-chemoproteomics workflow in ATRX^KO^ cells. **B.** Volcano plot of proteins competed upon cell treatment with 50 µM PP12. The dataset has been coloured with hits from the SL genetic screen performed by Segura-Bayona et al (in blue). N=3. **C/D.** Volcano plot of proteins competed upon cell treatment with expanded concentrations range: PP12 (C) PP734 and PP735 (D). The dataset reported is filtered for genetic hits. N=3. **E.** Percentage TYMS competition upon dosing with PP12/PP734/PP735. Significance threshold (50%) is noted with a dotted line. *P*-values 75 µM (PP12:PP734: 0.75, PP12:PP735: 0.96); *P*-values 50 µM (PP12:PP734: 0.15, PP12:PP735: 0.67); *P*-values 30 µM (PP12:PP734: 0.22, PP12:PP735: 0.60) – obtained with the Tukey’s multiple comparisons test on PRISM. N=3. **F.** 5-FU competition of PP1046 in ATRX^KO^ cells. In the top left quadrant of the volcano plot are reported proteins which have been competed >50%. N=3.

PP12 pre-treatment (4 h, 50 μM) reduced PP1046-mediated pulldown of 37 proteins (hit-calling ≥50% competition) (**Fig. 4b**). Among the most highly competed proteins were selenoproteins (SELENOK, SELENOS, SELENOH) and protein-disulfide reductase family members (TXNDC12, TXNDC17). Given the high intrinsic reactivity of selenocysteine and catalytic thiols in thiol–disulfide oxidoreductases, these were considered likely off-targets. To prioritize biologically relevant candidates, we cross-referenced competed proteins with a published ATRX-deficient genetic dropout dataset^8^. Filtering the chemoproteomics hits against known ATRX synthetic lethal genes yielded a single candidate: TYMS, a rate-determining enzyme in DNA synthesis^34^, which was robustly competed by PP12 in a concentration-dependent manner (**Fig. 4c**). (TXN was also strongly competed at high concentrations but was excluded from further analysis owing to the presence of hyperreactive cysteines common to thioredoxin family proteins^35^).

To further support TYMS as the phenotype-driving target, we repeated competitive profiling with the stereoprobes PP734 and PP735 (**Fig. 4d**). Both enantiomers produced strong, comparable competition for TYMS at 75, 50, and 30 µM (**Fig. 4d, 4e**), consistent with the lack of stereoselectivity observed in phenotypic assays.

### PP12 binds covalently to TYMS *in vitro*

TYMS (thymidylate synthase) is a rate-determining enzyme in the *de novo* synthesis of dTMP. Several cysteine residues have been implicated in TYMS function^36^, including the active-site cysteine C195^37^, which represents a potential ligandable site for a chloroacetamide-based reactive fragment such as PP12. TYMS catalyses the reductive transfer of a methylene group from the 5,10-methylene tetrahydrofolate (mTHF) cofactor to dUMP to form dTMP (**Fig. 5a**). During this process, C195 acts as a nucleophile, attacking the pyrimidine ring of dUMP at C6 to activate the C5 position for subsequent reaction with mTHF.

**Fig. 5.**
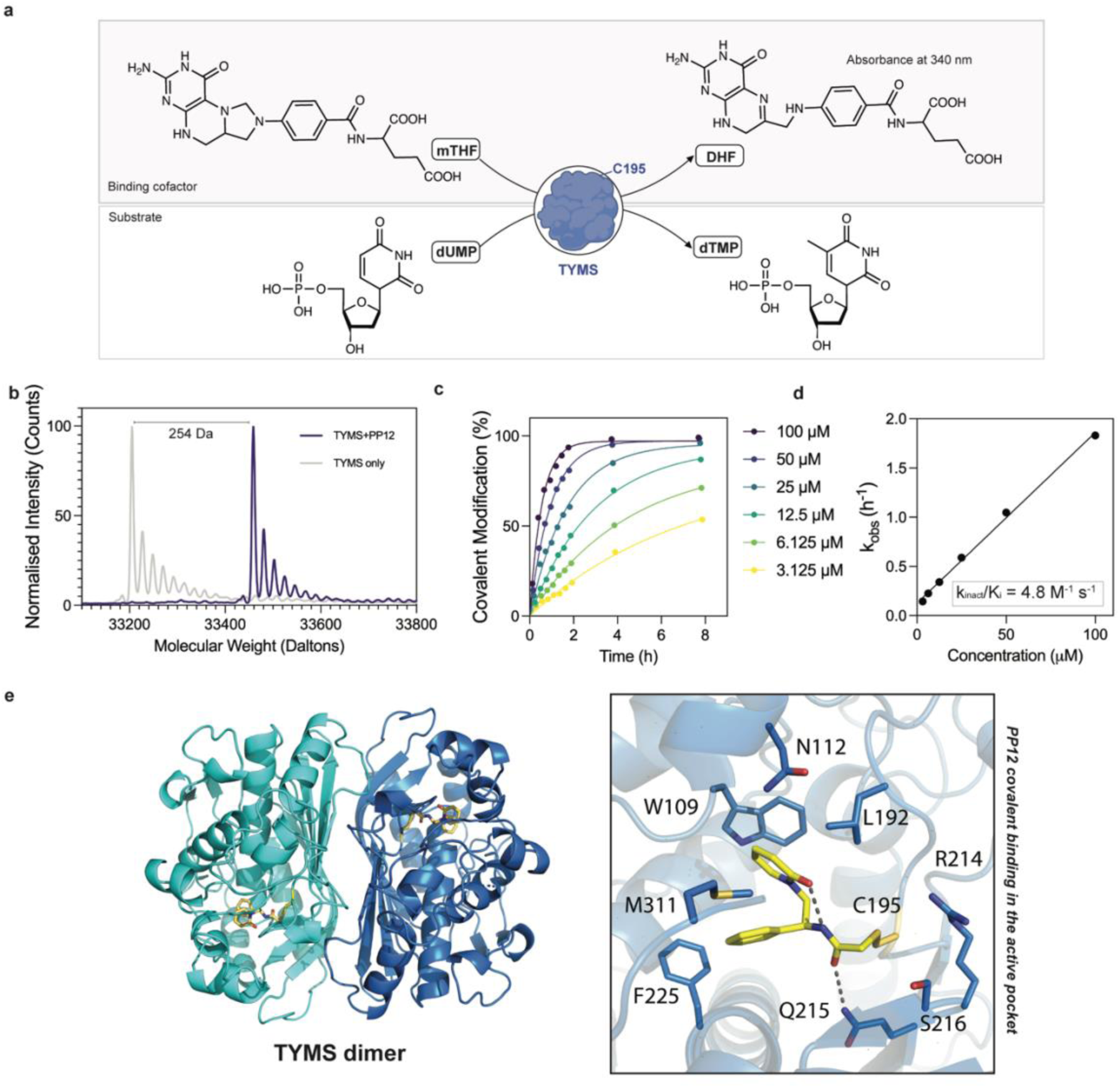
PP12/TYMS in vitro binding validation. **A.** TYMS mechanism of action. TYMS converts dUMP into dTMP by reducing the mTHF co-factor to DHF. **B.** TYMS-PP12 binding via intact protein LC-MS spectra. TYMS-only sample (in grey) compared to TYMS samples incubated with 100 µM PP12 (in blue). **C.** Time and concentration dependent TYMS-PP12 binding *via* intact protein LC-MS. **D.** TYMS-PP12 labelling k_inact_/K_i_. **E.** Crystal Structure of Homodimer of TYMS with covalently bound PP12 in the active site (LHS). Close-up of the C195-PP12 adduct with residues of the active site first shell within 5 Å of the inhibitor (RHS): hydrogen bonds shown as black dashes.

TYMS is an established cancer target that can be inhibited by several therapeutic agents. The antimetabolite 5-fluorouracil (5-FU), which is approved to treat cancers including solid carcinomas^38^, functions as a prodrug that is metabolised to 5-fluorodeoxyuridine monophosphate (FdUMP). FdUMP forms a stable ternary complex with TYMS and mTHF^39^, resulting in covalent inhibition of the active site through engagement of C195^40,41^.

To provide orthogonal evidence for TYMS as the PP12 target, we repeated the competitive click-chemoproteomics using 50 µM 5-FU prodrug (Sigma). We reasoned that if 5-FU and PP1046 engage the same site, 5-FU pre-treatment would reduce TYMS enrichment by PP1046. Strikingly, TYMS was the only protein competed >90% (**Fig. 4f**), consistent with PP12/PP1046 and FdUMP occupying the same active-site region.

We next measured PP12 covalent modification of recombinant TYMS *in vitro* by intact-protein LC–MS. PP12 treatment (100 – 3.125 μM) resulted in time- and concentration-dependent labelling of TYMS (**Fig. 5b**, **Fig. 5c**) and fitting yielded a *k*_inact_/*K*_i_ value of 4.8 M^-1^ s^-1^ (**Fig. 5d**)^42^.

Finally, we asked whether the covalent binding occurred at the active site *via* crystallography approaches. The structure of the PP12-bound TYMS_26-313_ was solved at 2.4 Å by molecular replacement using the available coordinates of the human TYMS in complex with dUMP (PDB ID 6QXH^40^) with the ligand removed (**Fig. 5e**, Supplementary Table 1). Only one protomer of the constitutive dimer was present in the crystallographic asymmetric unit with the dimeric enzyme reconstructed by symmetry operations (**Fig. 5e**, LHS).

Similarly to previously reported complexes presenting dUMP and antifolates inhibitors^40,43^, the catalytic loop (residues 181–197) of the PP12-bound TYMS_26-313_ structure was observed to adopt an active conformation. Small deviations from ligand-containing literature structures^40^ were observed in the conformation of loop 46-53 due to crystal contacts.

Well-defined positive electron density was present in the active site in continuity with the catalytic C195 sulfur atom, confirming the presence of the adduct formed by PP12 with the cysteine sidechain (**Fig. 5e**). Bound PP12 shows the pyridone and the phenyl rings in a reciprocal stacking position and its positioning in the pocket would hamper the binding of both dUMP and mTHF. Several close contacts with the surrounding residues are observed **(Fig. 5e)** including edge to face *π*-stacking interactions between the phenyl ring with F225 and the pyridone with W109, hydrophobic interaction between L192 and the pyridone, in addition to H-bonding of the carbonyl with NH2 of Q214 (3.2 Å). An internal H-bond between PP12 amidic NH and the pyridone O stabilises the orientation of the ring (2.6 Å).

### dTMP cellular levels are responsible for SL in ATRX-deficient background

Having established that PP12 binds to the catalytic cysteine of TYMS, we investigated how TYMS inhibition could drive synthetic lethality in an ATRX-deficient context. We first sought to establish whether PP12 perturbs the canonical role of TYMS in dTMP synthesis (**Fig. 6a**). To this end, we adapted an established TYMS spectrophotometric activity assay (*Wahba et al.*)^44^ monitoring mTHF conversion to DHF by absorbance at 340 nm (**Fig. 5a**). Enzyme activity was measured in the presence of PP12 (1000–1.9 μM) or DMSO and quantified using four-parameter competitive inhibition kinetics, revealing a marked reduction in activity upon PP12 treatment (**Fig. 6b**). Notably, apparent IC_50_ values were dependent on PP12 pre-incubation time, shifting from 119 ± 15 μM (10 min) to 18 ± 2 μM (30 min).

**Fig. 6.**
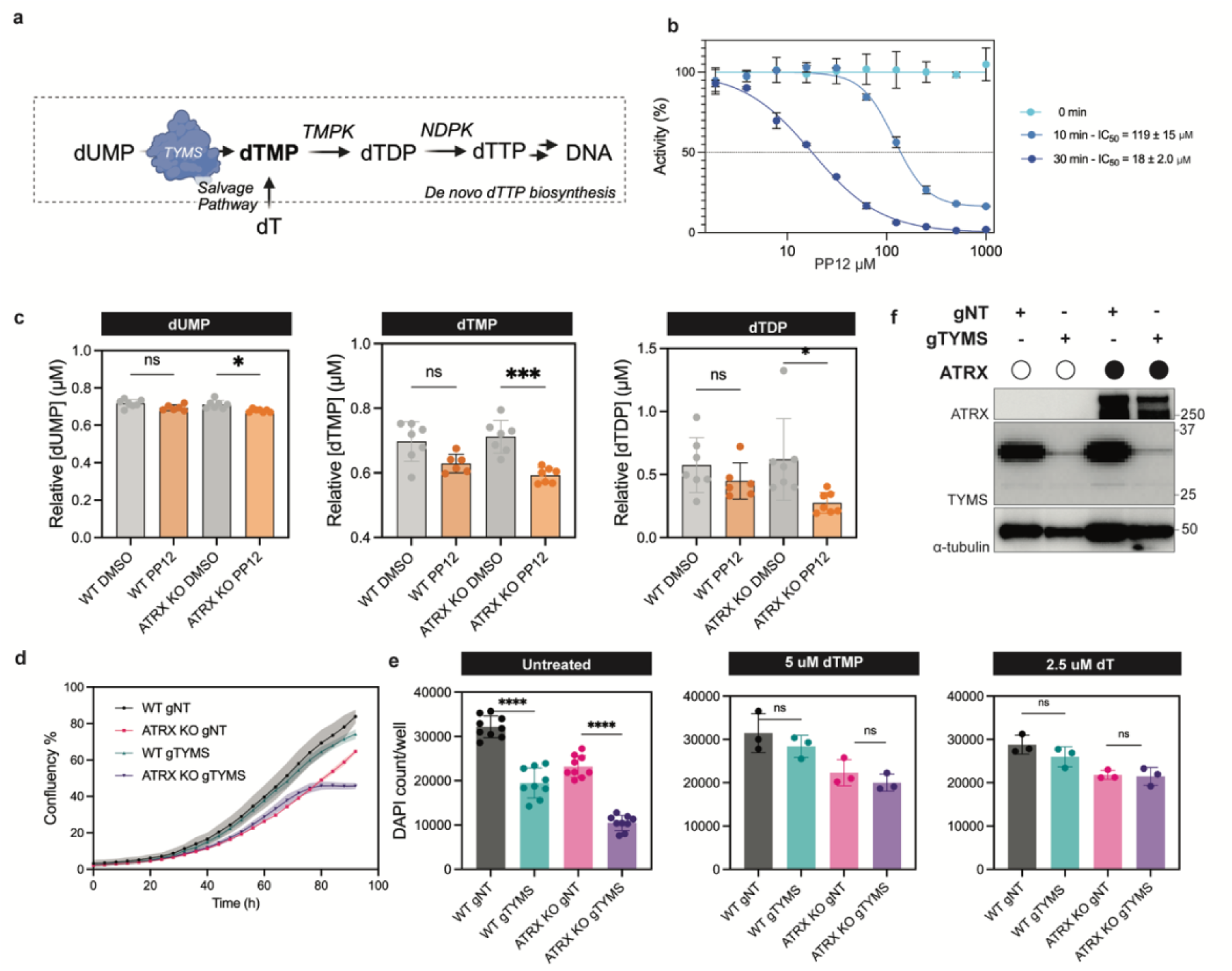
PP12 affects dTMP metabolite levels in cells with an effect on the SL phenotype. **A.** dTMP *de novo* synthetic route. Figure is adapted from Ref^34^. **B.** TYMS Kinetic Inhibition *in vitro* monitored via absorbance of DHF. N=2. **C.** Targeted LC-MS analysis of dTMP, dUMP and dTDP metabolites levels upon treatment. N=6. *P*-values dUMP (WT: 0.12, ATRX KO: 0.01), dTMP (WT: 0.05, ATRX KO: < .001), dTDP (WT: 0.72, ATRX KO: 0.03 – obtained with the ordinary one-way ANOVA statistical test on PRISM. **D.** Viability of eHAP iCas9 ATRX^KO^ TYMS^KO^ (gTYMS), eHAP iCas9 WT TYMS^KO^ (gTYMS), eHAP iCas9 ATRX^KO^ (gNT), eHAP iCas9 WT (gNT) monitored via Incucyte readout. N=3. **E.** Viability of eHAP iCas9 ATRX^KO^ TYMS^KO^ (gTYMS), eHAP iCas9 WT TYMS^KO^ (gTYMS), eHAP iCas9 ATRX^KO^ (gNT), eHAP iCas9 WT (gNT) measured as DAPI nuclear count. *P*-values Untreated Conditions (WT gNT – WT gTYMS: < .001, ATRX KO gNT – ATRX gTYMS: < .001), 5 µM dTMP (WT gNT – WT gTYMS: 0.44, ATRX KO gNT – ATRX gTYMS: 0.66), 2.5 µM dT (WT gNT – WT gTYMS: 0.53, ATRX KO gNT – ATRX gTYMS: 0.10) – obtained with the unpaired t-test statistical test on PRISM. **F.** TYMS levels monitored *via* western blot analysis upon 4 days of induction.

We next assessed whether PP12 reduced dTMP levels in cells. Accordingly, we performed targeted metabolomics following treatment of WT and ATRX^KO^ iCAS9 eHAP cells with PP12 (33 μM) for 24 h. dTMP levels were reduced relative to DMSO controls in both cell lines (**Fig. 6c**), indicating that PP12 perturbs TYMS function in cells. Although dTMP reduction was marginally more pronounced in ATRX^KO^ cells, we attribute this to reduced viability under treatment (**Fig. S5e**). A similar decrease was observed for dTDP (a downstream metabolite), whereas levels of the precursor dUMP were comparable between treated and untreated samples (**Fig. 6c**).

To test whether the ATRX/TYMS SL interaction could be rescued by restoring thymidylate pools, we explored supplementation with pathway products. Doxycycline-inducible Cas9 cell lines were generated, expressing sgRNAs targeting TYMS or a non-targeting control (gNT). Knockout efficiency was monitored by western blot and showed a substantial reduction in TYMS levels by day 4 after Cas9 induction (**Fig. 6f**). Cell growth was tracked by Incucyte imaging over four days following induction, after which cells were fixed, stained with DAPI, and quantified using the Opera Phoenix Imaging system. Consistent with a synthetic lethal interaction, growth of ATRX^KO^ TYMS^KO^ cells was strongly impaired upon induction, whereas WT TYMS^KO^ cells were less affected (**Fig. 6d**). Importantly, supplementation of the culture media with TYMS pathway products (dTMP, dTTP, or dT) fully rescued the viability defect in TYMS^KO^ cells (**Fig. 6e**).

### PP12 reveals global replication stress as a determinant of ATRX/TYMS synthetic lethality

ATRX is a SNF2 family helicase/ATPase with a conserved ATPase domain^45^ and additional domains mediating interactions with chromatin and replication factors, including a PCNA-interacting protein (PIP) motif^46,47^ (**Fig. 7a**). Previous work has shown that ATRX synthetic lethality can arise through distinct mechanisms linked to global DNA replication and/or telomere maintenance^8^ and that these functions can be genetically separated through ATRX PIP-box and ATPase activities (**Fig. 7a**).

**Fig. 7.**
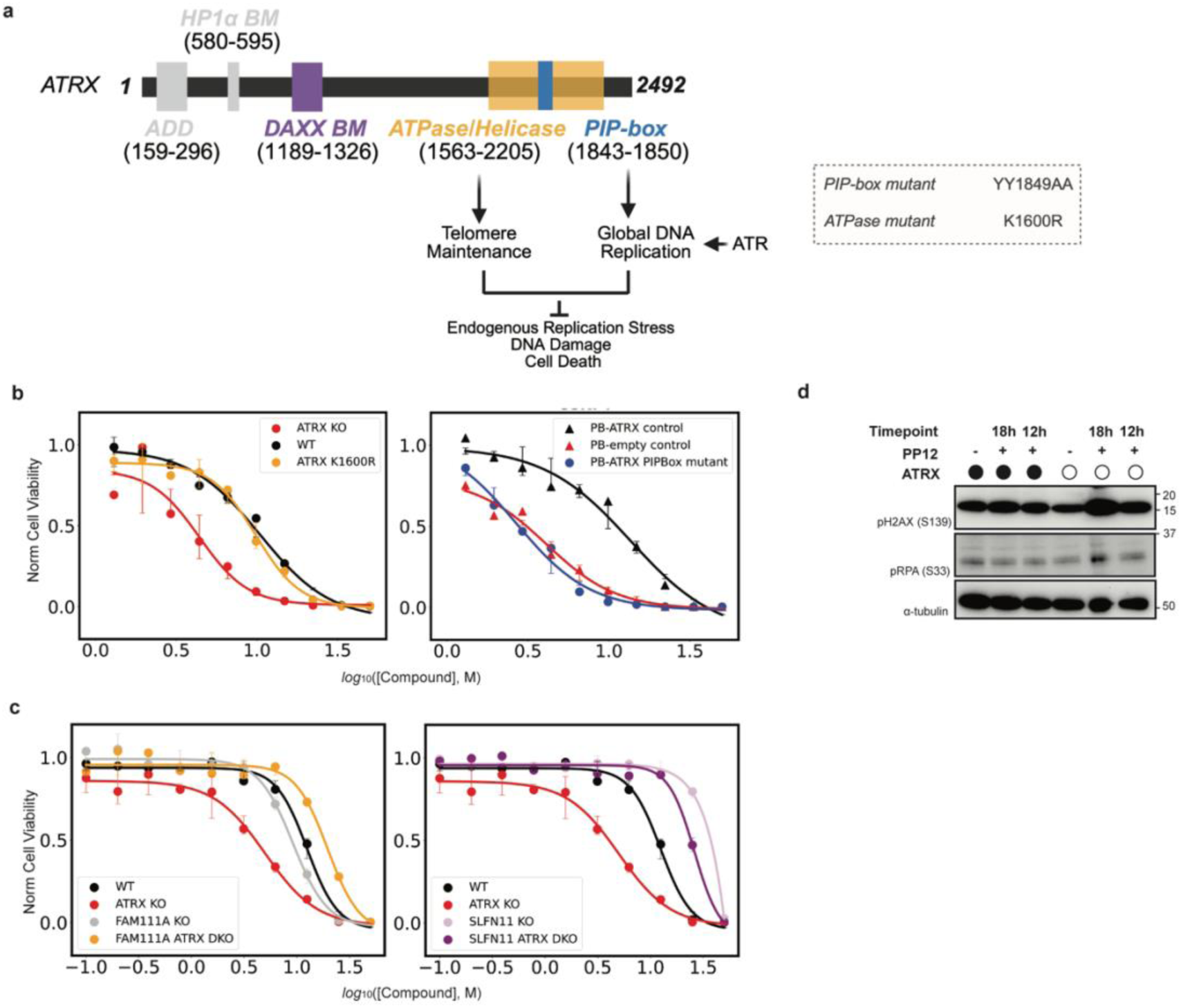
PP12 as a tool to further understand the role of TYMS in ATRX biology. **A.** ATRX protein key domains. Figure adapted from *Segura-Bayona et al*^40^ **B.** PP12 concentration-response behaviour in ATRX-ATPase mutant and PIP-Box domain mutant cell lines monitored *via* CTG. N=3. **C.** Viability rescue of SL upon SLFN11^KO^ (RHS) or FAM111A^KO^ (LHS) monitored via CTG. N=3. **D.** Western blot Analysis of DNA replication stress markers pRPA and γH2AX upon treatment.

We therefore questioned whether PP12-induced synthetic lethality depends on specific ATRX functions. We treated a panel of cell lines expressing WT ATRX, or ATRX mutations in the PIP-box (ATRX^KO^ complemented with PiggyBac-ATRX^PIPmut^), or ATPase domain (ATRX^KO^ with ATRX^K1600R^ knock-in) with PP12^8^. ATPase domain mutation did not significantly alter viability relative to the WT ATRX controls, whereas PIP-box mutation phenocopied ATRX^KO^ cells (**Fig. 7b**), indicating that PP12 sensitivity depends on ATRX functions associated with DNA replication. Consistent with this model, PP12 synthetic lethality was independent of ATRX/DAXX-related activity (**Fig. S5d**).

*Segura-Bayona et al.* also reported that synthetic lethality in an ATRX-deficient background which is associated with global DNA replication stress can be rescued by loss of FAM111A^8^. FAM111A is a protease that associates with active replication forks^48^ and accumulates on chromatin in response to replication stress in ATRX-deficient cells. By showing resistance to PP12 in ATRX^KO^ FAM111A^KO^ cells, we were able to support a replication-stress related mechanism for the ATRX/TYMS SL interaction (**Fig. 7c**).

Loss of SLFN11, a tRNA nuclease and putative helicase that is epigenetically silenced in ∼50% of tumours^49^, has been shown to protect ATRX-deficient cells from apoptosis and to rescue ATRX synthetic lethality with ATR inhibitors^50^. Accordingly, PP12 treatment of ATRX SLFN11^DKO^ cells did not reduce viability relative to the eHAP WT control (**Fig. 7c**). This result further supports the specificity of the pathway by which PP12 induces synthetic lethality in an ATRX-deficient background.

Finally, we monitored replication stress and DNA damage markers to assess whether PP12 induces global replication stress selectively in ATRX-deficient cells. PP12 treatment increased phosphorylation of RPA at S33 (pRPA S33), a marker of replicative stress^51^, and increased γH2AX, a marker of DNA double-strand breaks^52^, in eHAP ATRX^KO^ cells but not in WT controls (**Fig. 7d**).

## DISCUSSION

In this work, we describe a phenotypic screening approach for covalent fragment libraries that exploits synthetic lethality in an ATRX-deficient setting. Although covalent compound libraries have previously been screened directly in cells^16,25^, identifying interactions with meaningful biologically activity remains challenging. By starting from a disease-relevant cellular phenotype, this approach prioritises compounds with functional effects and enables streamlined progression from phenotype to target and mechanism.

ATRX is among the most frequently mutated proteins in cancer and has been reported to exhibit synthetic lethality with hundreds of partners^8^, making ATRX-deficiency a tractable phenotype for chemical interrogation. Tractable examples of synthetic lethal interactions in ATRX-deficient settings include PARP inhibition^10^, WEE1 inhibition^53^, ATM inhibition^54^, and G4 stabilisation^55^.

Using viability screening isogenic ATRX WT/KO cell lines, we identified a chloroacetamide fragment (PP12) that reproducibly and selectively reduces viability in ATRX-deficient cells (**Fig. 2**). To deconvolute the molecular targets responsible for this phenotype, we implemented a competitive click-chemoproteomics workflow. As anticipated for a chloroacetamide fragment, PP12 engaged multiple proteins in live cells^56^, highlighting the difficulty of assigning phenotypic relevance based on chemoproteomic screening alone (**Fig. 3**). By subsequent integration of data from a published genome-wide dropout screen^8^, we converged on a single candidate target: TYMS (**Fig. 4c**). Orthogonal competitive profiling with 5-fluorouracil (a known TYMS inhibitor) further supported engagement of the TYMS active-site region. Crystallography and intact-protein LC-MS revealed covalent engagement of C195, the active-site cysteine of TYMS (**Fig. 5**).

TYMS is a validated cancer target, with widespread clinical use of TYMS-directed antimetabolites; however their utility is limited by toxicity, resistance and variable response^57–59,38,60^. Our findings support TYMS as an ATRX synthetic-lethal partner, suggesting that ATRX-deficient contexts may confer heightened sensitivity to TYMS inhibition.

We validated the ATRX–TYMS synthetic lethal interaction by generating an inducible ATRX^KO^ TYMS^KO^ cell line and assessing viability under defined growth conditions. Induction of TYMS knockout substantially impaired viability in the ATRX^KO^ background, supporting the synthetic lethal interaction. Moreover, supplementation with dTMP, dT, or dTTP fully rescued the viability defect, supporting a model in which reduced thymidylate availability is directly linked to the heightened toxicity observed upon TYMS loss in ATRX-deficient cells (**Fig. 6**).

Finally, we used PP12 to probe the mechanistic basis of ATRX/TYMS synthetic lethality in the context of DNA damage. Previous studies have demonstrated that ATRX deficiency can be synthetic lethal with different partners through distinct ATRX functions^8^: the PIP motif is linked to regulation of global DNA replication, whereas the ATPase domain contributes to suppression of single-stranded telomeric DNA. By using ATRX separation-of-function mutants and genetic modifiers of replication stress, we provide evidence that the PP12/TYMS synthetic-lethal interaction depends on the ATRX PIP-box function and is rescued by loss of FAM111A and SLFN11. These observations are consistent with TYMS inhibition driving loss of viability in ATRX-deficient cells through global replication stress (**Fig. 7**).

In summary, these results demonstrate that a ‘phenotype-first’ covalent screening approach can identify tool molecules with disease-relevant synthetic lethal activity, enable target deconvolution, and support mechanistic validation in cells. This approach helps address two persistent challenges in early-stage drug discovery: identification of actionable targets in a disease-relevant context and rapidly generating small molecules that engage those targets in cells as starting points for lead discovery. By beginning with a cellular phenotype, we avoid common hurdles of chemoproteomics-first approaches, including the difficulty of assigning functional consequences to target engagement. A challenge of this approach is that reactive fragment libraries must strike a balance between proteome coverage, cellular potency, and sufficient selectivity to enable phenotype modulation, while also allowing polypharmacology to be identified and disentangled. Tuning electrophile reactivity and fragment size/complexity is central to this trade-off: less reactive moieties (e.g., acrylamides) may yield cleaner, more selective hits, but can shift screening requirements toward larger libraries (or higher concentrations/longer incubations) to achieve comparable coverage and phenotype discovery. As chemoproteomics datasets on proteome-wide engagement of covalent libraries grow, machine-learning approaches will likely be impactful in guiding the designing of libraries that maximise the potential for phenotype modulation for a given library size^61,62^.

Importantly, in this study the integration of chemoproteomic competition with genetic dropout data was a major asset in resolving phenotypically relevant targets from a background of multiple competed proteins; in settings where comparable genetic datasets are unavailable, target assignment may require alternative triangulation strategies, including causal-biology computational approaches and more chemically controlled deconvolution (e.g., matched enantiopairs, close non-hit analogues, or rapid, iterative hit expansion to build structure–phenotype and structure–engagement relationships, potentially via direct-to-biology-style workflows)^63,64^.

Although TYMS is a well-established cancer target with clinically used inhibitors, our work validates its role as a synthetic lethal partner with ATRX and points to new opportunities to exploit this vulnerability. More broadly, these findings show that a phenotype-first, target- and partner-agnostic platform can connect covalent fragment phenotypes to molecular targets and mechanisms, enabling discovery of new synthetic lethal pairs and chemical starting points for future therapeutic development.

## METHODS

### Cell lines information and culture conditions

eHAP iCas9 WT c3.22, eHAP iCas9 ATRX^KO^ g3, eHAP iCas9 ATRX^KO^ g4, eHAP iCAS9 ATRX^(K1000R)^, eHAP iCAS9 ATRX^KO^ (PB-ATRX^PIP-box^), eHAP iCAS9 ATRX^KO^ (PB-empty control), eHAP iCAS9 ATRX^KO^ (PB-ATRX control), eHAP iCAS9 ATRX^KO^ (PB-ATRX^DAXXmutant^), eHAP iCAS9 SLFN11^KO^, eHAP iCAS9 FAM111A^KO^, eHAP iCAS9 ATRX ^KO^ SLFN11^KO^, eHAP iCAS9 ATRX ^KO^ FAM111A^KO^ were maintained by passaging upon reaching 70-80% confluency in Iscove’s modified Dulbecco’s medium IMDM (Gibco, 21980-032) 10% Tet-Free FBS (South America origin, Fetal bovine serum, 0.2 µm sterile filtered, 500ml PAN Biotech, P30-3602) and 1% PenStrep (Gibco, 15140-122) media conditions (growth conditions = 37C, 5% CO2). Cell lines were generated as in *Segura-Bayona et al*^50^.

NCI-H460 ATRX-KO c10 and NCI-H460 iCas9 C19 were maintained by passaging upon reaching 70/80% confluency in Roswell Park Memorial Institute RPMI Medium 1640 (Gibco, 21875-034), 10% Tet-Free FBS (South America origin, Fetal bovine serum, 0.2 µm sterile filtered, 500ml PAN Biotech, P30-3602) and 1% PenStrep (Gibco, 15140-122) media conditions (growth conditions = 37C, 5% CO2). Cell lines were generated as in *Stanage et al*^65^ and *Segura-Bayona et al*^50^.

eHAP iCas9 WT gNT, eHAP iCas9 ATRX^KO^ gNT, eHAP iCas9 WT gTYMS, eHAP iCas9 ATRX^KO^ gTYMS, were maintained by passaging upon reaching 70-80% confluency in in Iscove’s modified Dulbecco’s medium (Gibco, 21980-032), 10% Tet-Free FBS (South America origin, Fetal bovine serum, 0.2 µm sterile filtered, 500ml PAN Biotech, P30-3602) and 1% PenStrep (Gibco, 15140-122) media conditions (growth conditions = 37C, 5% CO2). Cells are selected with 0.4 µg/µL puromycin for two days prior to experiments and knockout is induced by adding 1 µg/µL of doxycycline.

### High Throughput Screening and Data Acquisition

10 µL of media containing the reactive fragments at concentrations of 500, 100 and 10 µM (or DMSO control) were added to individual wells in 384-well plates Greiner µclear Cat: 781091. eHAP iCas9 WT and ATRX^KO^ g3 cells were seeded at 200 cells/well in 90 µL of media, for final concentrations of reactive fragments of 50, 10 and 1 µM (or equivalent DMSO volume). Confluency of the WT DMSO control was monitored with the IncuCyte S3 System (Sartorius) and plates were fixed (4% formaldehyde) and stained with DAPI (Merk D9542) (1:10000) after 4 days. Plates were imaged by CellInsightCX7 (Thermo Scientific) using 10x, 9 fields to cover the whole well. The raw data is analysed through SAS web analysis pipeline using web CellHTS2^66^.

### High Throughput Screening Data Analysis and hits selection

Cell viability for each condition is assessed by calculating the median of the nuclei count normalized by the relative DMSO control (and the associated MAD value) across three replicates (note that data has been previously normalized by the Z-score correction as highlighted in the python scripts provided in the supplementary materials). The three-point concentration-response curves (for each compound in each cell line) are fit to a restrained logistic regression on each 3-point nuclei count curve to allow the curve midpoint (pEC50) and area under the curve (AUC) of the logistic function to be calculated. The difference between these values for a given compound in both cell lines is additionally calculated as a measure of the SL phenotype. AUC errors are calculated by Monte Carlo error propagation (10,000 iterations). The ΔAUC is estimated to be 0 for the compounds where calculated pEC50 values lay outside of the boundary of the concentration range examined. **Fig S1a** reports the full range of compounds including those which were filtered out. Hits are identified as those compounds that show an *absolute* ΔAUC (logistic regression values) ≥ mean ± 2*stdev. Compounds are selected for further validation based on:

- The strength of the phenotype observed.
- Structural similarity of a compound to the identified hits.

Structural similarity is assessed by calculating Tanimoto similarity scores between Morgan fingerprints, followed by hierarchical clustering using the Ward algorithm^67^ with RDKit (https://www.rdkit.org). Compounds that have a pairwise Tanimoto similarity score of ≥0.4 to one of the hits identified by hit calling is selected for further analysis. The dendrogram and selected hits are reported in **Fig. S1b**.

### Hits fragments validation *via* concentration-response Data Acquisition

The experimental protocol can be found in the previous high throughput screening section. A varying concentration range is used for acrylamides reactive moieties (1-50 μM (12 concentrations) or 1-100 μM (7 concentrations)) and chloroacetamides reactive species (1-50 μM or 0.25-20 μM (12 concentrations) to obtain concentration-response curves.

### Hits fragments validation *via* concentration-response Data Analysis and hits selection

Nuclei count is normalised with respect to in-plate untreated cells and the relative DMSO-control. Data is fit to a logistic regression and ΔpEC50 is extrapolated from the fitting. The top hits are selected according to their ΔpEC50 value and quality of the fitting by inspecting the resulting curves.

### CellTitre-Glo viability assay experimental and data analysis

Cell lines are seeded at 200 cells/well (eHAPs) or 400 cells/well (NCI-H460) in 200 µL of media in a 96-well plate (Greiner 655098) (day 0). Cells are subsequently dosed on day 1 with reactive fragments (or DMSO control) of interest to obtain a range of concentrations. Plates are kept at 37C, 5% CO2 for an additional 5 days. Old media is then discarded, substituted with 1X CellTiter-Glo® One Solution Assay and incubated at RT for 10 minutes with shaking. Luminescence reading is performed with the ClarioStar^Plus^ Plate reader.

Data were normalised for untreated conditions and relative DMSO controls for each fragment and averaged across biological replicates (median). Details on fitting, pEC50 calculations and statistical relevance can be found in the relevant python scripts and Prism files associated with the manuscript.

### Mass spectrometry general

The Evosep One was coupled online to a hybrid (trapped ion mobility spectrometry) TIMS quadrupole TOF (time of flight) mass spectrometer (Bruker timsTOF Pro 2) via a captive spray nano-electrospray ion source. Samples were analysed with in DIA-PASEF mode using an ion mobility range from 1/K0 = 1.6 to 0.6 Vs cm^-2^. Data was acquired using variable ion mobility windows and variable mass windows, between a mass range of 262.18–1398.68 *m*/*z*^68^. The data acquisition method was optimised using pyDIAD^69^.

Raw mass spectrometry data files were analysed using Spectronaut (v18.7 or v19.1) with directDIA. The following search parameters were used for directDIA: peptide lengths of 7–52 amino acids with up to two miscleavages were permitted, with fixed modification of carbamidomethyl (C) and variable modifications: acetyl (Protein N-term), oxidation (M). All searches were performed against two FASTA files that contained the canonical UniProt human protein sequences^68^ and common contaminants.

### Competitive click chemoproteomics experimental

eHAP ATRX^KO^ g3 cells were seeded at 2M cells/dish in 10 cm dishes and left to attach overnight. The media was subsequently replaced with 3 mL of fresh media containing either competitor (at 75, 50, 30 or 15 μM) or DMSO vehicle for 4 hours. DMSO concentration = 0.1%. Then the alkyne probe (50 μM) (or DMSO) was added for 1 h. Pellets were then collected by centrifugation, washes with PBS and snap frozen on dry ice for −80 ℃ overnight storage.

### Competitive click chemoproteomic profiling

#### Acetylation of NeutrAvidin™ agarose beads

NeutrAvidin™ agarose, high-capacity beads (Thermo Fisher, #PIER29204) were centrifuged (2,000 rcf, 2 mins) and supernatant was removed. The beads were washed three times with PBS buffer, before PBS (9 mL) and Sulfo-NHS-Acetate (200 mM made up fresh in 965 μL anhydrous DMSO Thermo Fisher #26777) were added. The beads were incubated for 30 mins at room temperature on a falcon tube roller, then centrifuged (2,000 rcf, 2 mins). The supernatant was removed and the incubation step with freshly made up Sulfo-NHS-Acetate was repeated. The reaction was quenched by adding Tris (1 M, pH 7.5, 2 mL). The beads were centrifuged (2,000 rcf, 2 mins) and supernatant was removed, washed once with PBS, then twice with 20% EtOH. EtOH (20%, 10 mL) was added, and the beads were stored at 4℃ until required.

#### Cell lysis

The cells were lysed over ice with 250 μL of freshly made lysis buffer for 45 mins (SDS 0.1%, 1% IGEPAL, 0.5% Na-deoxycholate, 150 mM NaCl, 50 mM HEPES pH 8.0, EDTA-free protease inhibitor cocktail, 1:1000 benzonase). The protein content of each sample was determined by BCA assay, 400 μg of lysate was taken forward (total volume adjusted to 376 μL with lysis buffer, 1.06 μg/μL).

#### Click reaction

24 μL of premixed click mixture was added to each sample (final concentrations biotin-PEG3-azide 165 μM, 0.65 mM CuSO4, 0.65 mM BTTAA^33^, 1.1 mM aminoguanidine, 1.1 mM ascorbic acid, final protein concentration 1.0 μg/μL) and samples shaken (1,000 rpm) at room temperature for 1 hour. The click reaction was quenched by the addition of EDTA (8 µL, 0.5 M, 10 mM final concentration). Proteins were precipitated with the addition of 1.6 mL of ice-cold acetone and stored at −20℃ overnight. The resulting protein pellets were washed twice with ice-cold 80% acetone (vigorous sonication was required to fully break the protein pellet each wash). The air-dried pellets were dissolved in 0.2% SDS 50 mM HEPES pH 8.0 by vortexing and sonicating.

#### Bead enrichment and digestion

Samples were incubated with previously prepared acetylated NeutrAvidin™ agarose resin (50 μL of bead slurry) pre-washed three times with 0.2% SDS in 50 mM HEPES pH 8.0) on a sealed combinatorial microlute plate for 2 hours. The plate was centrifuged (700 rcf, 1 min). The beads were washed (3 × 850 μL) with 0.1% SDS in 50 mM HEPES pH 8, 4 M urea in 50 mM HEPES pH 8.0) and HEPES (50 mM pH 8.0). The enriched proteins were reduced and alkylated on bead (5 mM TCEP, 15 mM chloroacetamide in 50 mM HEPES pH 8.0) for 30 mins, then washed once with 850 μL of 50 mM HEPES pH 8.0. The proteins were then digested at room temperature overnight by the addition of 60 μL of freshly made digestion buffer (0.003 μg/μL LysC in 50 mM HEPES pH 8.0).

#### Peptide elution and further digestion

The plate was placed on a clean collection plate and centrifuged to collect the peptides. An in-solution trypsin digest was performed (30 μL of 0.003 μg/μL trypsin in 50 mM HEPES pH 8.0) and the plate was incubated at 37℃ for 4 hours. The trypsin was inactivated by acidification (10 μL of 10% formic acid).

#### LC-MS/MS analysis

Samples were loaded onto Evotips following conditioning and preparation recommended by the manufacturer, then injected onto the Bruker timsTOF Pro 2 mass spectrometer coupled with an Evosep One LC system using 60 samples per day method.

#### Data analysis

See Mass spectrometry: general. Data was then further processed in Perseus^70^. Intensity values were then log2 transformed and missing values imputed from normal distribution (width = 0.3, down shift = 1.8, separately for each column). Log2 fold change and -log_10_(*p*-value) were then calculated between different conditions using T-test.

#### Spectra normalisation

Data was normalised prior to plotting with max/min normalisation. Refer to supplementary materials for additional information.

### TYMS protein expression and purification

TYMS_26-313 synthetic gene: cctctttcagggacccggtccgccgcacggggagctgcagtacctggggcagatccaacacatcctccgctgcggcgtcaggaaggacgaccgcacgggcaccggcaccctgtcggtattcggcatgcaggcgcgctacagcctgagagatgaattccctctgctgacaaccaaacgtgtgttctggaagggtgttttggaggagttgctgtggtttatcaagggatccacaaatgctaaagagctgtcttccaagggagtgaaaatctgggatgccaatggatcccgagactttttggacagcctgggattctccaccagagaagaaggggacttgggcccagtttatggcttccagtggaggcattttggggcagaatacagagatatggaatcagattattcaggacagggagttgaccaactgcaaagagtgattgacaccatcaaaaccaaccctgacgacagaagaatcatcatgtgcgcttggaatccaagagatcttcctctgatggcgctgcctccatgccatgccctctgccagttctatgtggtgaacagtgagctgtcctgccagctgtaccagagatcgggagacatgggcctcggtgtgcctttcaacatcgccagctacgccctgctcacgtacatgattgcgcacatcacgggcctgaagccaggtgactttatacacactttgggagatgcacatatttacctgaatcacatcgagccactgaaaattcagcttcagcgagaacccagacctttcccaaagctcaggattcttcgaaaagttgagaaaattgatgacttcaaagctgaagactttcagattgaagggtacaatccgcatccaactattaaaatggaaatggctgtttaataacgctctggtgccacgcgg Recombinant human TYMS_26-313_, a truncated construct lacking the first N-terminal 25 aa which are unstructured and not located in the density of the human full-length structure, was expressed in E. coli BL21 cells using a pET49 plasmid with an amino-terminal cleavable His-tag. After cells harvesting, resuspension in buffer A (50 mM Hepes pH 7.5, 300 mM NaCl, 20 mM imidazole, 0.5 mM TCEP), lysis by sonication and centrifugation, the protein was purified by Ni-NTA affinity chromatography using column equilibration buffer A. After loading of the lysate, followed by wash with buffer A and buffer B (50 mM Hepes pH 7.5, 1 mM NaCl, 20 mM imidazole, 0.5 mM TCEP) the protein was eluted with buffer C (50 mM Hepes pH 7.5, 300 mM NaCl, 300 mM imidazole, 0.5 mM TCEP) After cleavage of the His-tag by 3C-protease overnight at 4 °C and concentration, a final purification step was performed by gel filtration using a 26/60 Sephadex column equilibrated with the storage buffer 50 mM Hepes pH 7.5, 50 mM NaCl, 2 mM TCEP. The protein was then concentrated to 20.2 mg/mL.

SEC chromatogram is reported in **Fig. S3c**. SDS-PAGE gel reported in Fig **S3d**.

### Covalent in vitro inhibition of TYMS

TYMS in vitro activity assay was adapted from previously reported literature^44^. Solutions of TYMS (150 μM), mTHF (150 μM), dUMP (200 μM) and PP12 (1000 – 1.9 μM) were prepared in HEPES pH 7.5 (50 mM) and 25 MgCl2 (25 mM) buffer with the following steps: Buffer, TYMS and mTHF were left to equilibrate at 25C for five minutes prior to PP12 addition. A positive control (no PP12) and a negative control (no TYMS) were subjected to the same conditions. The solutions were then left to incubate at 25C for 0, 10 or 30 minutes prior to substrate addition. dUMP was added to start the enzymatic reaction – which was recorded over time with absorbance at 340 nm. Fitting of the curves was performed in PRISM Graphpad.

### Kinetics of covalent modification by intact protein LC-MS

A stock of TYMS (0.5 μM) was prepared in 25 mM HEPES, 50 mM NaCl (pH 7.4). Compound of interest (PP12) was serially diluted across a 384 well-plate. An aliquot of the TYMS stock (10 µL) was dispensed to all wells. The plate was sealed, centrifuged (1000 rpm, 1 min, 4 °C) and then sampled immediately for IP-LC/MS analysis. Intact protein masses were recorded by LC-MS using an Agilent G6230 ToF Accurate Mass Series mass spectrometer, interfaced with an Agilent 1290 series liquid chromatography and sample handling system. Spectra were processed using Agilent MassHunter BioConfirm Software 10.0 with the Maximum Entropy method employed. The TIC was extracted (region containing protein) and the summed scans were deconvoluted (using a maximum entropy algorithm) over a *m*/*z* range (900 – 2,000) with the expected mass range (30,000 – 35,000).

### Agilent G6230 ToF

The protein sample was injected using an Agilent 1290 series AutoSampler (Model No. G7167B) with a 7 µL injection volume and maintained at a temperature of 20 °C. Chromatography was carried out on an Agilent Bio-HPLC polymeric reverse-phase (PLRP-S) (1000 Å, 5 µm × 50 mm × 1.0 mm, PL1312-1502) column at 70 °C. Using an Agilent 1200 series binary pump system (Model No. G6230B) the sample was eluted at 0.5 mL/min using a gradient system from solvent A (water, 0.2% formic acid (*v*/*v*)) to solvent B (acetonitrile, 0.2% formic acid (*v*/*v*)) according to the following conditions:

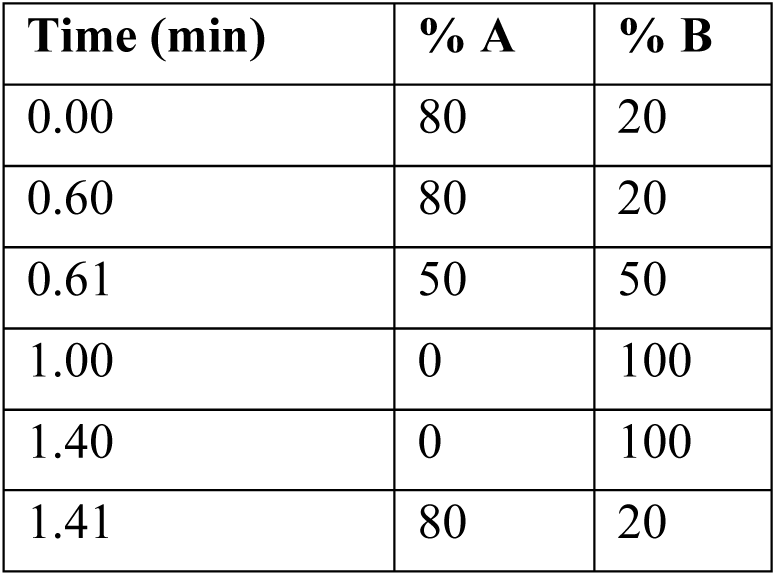

| Time (min) | % A | % B |
| --- | --- | --- |
| 0.00 | 80 | 20 |
| 0.60 | 80 | 20 |
| 0.61 | 50 | 50 |
| 1.00 | 0 | 100 |
| 1.40 | 0 | 100 |
| 1.41 | 80 | 20 |

The eluent was injected directly into an Agilent ToF mass spectrometer (Model No. G6230B) using a dual AJS ESI source and scanning between 600–3200 Da with a scan rate of 1.20 s in positive mode. The following MS parameters were used: capillary voltage limit 4000; desolvation temperature 350 °C; drying gas flow 10.0 L/min. Data acquisition was carried out in 2 GHz Extended Dynamic range mode.

The deconvoluted spectra were exported as csv files and analysed using R Studio software v3.5.1 to generate csv and pdf files of the spectra. Obtained IP-LC/MS spectra were visualised and analysed according to previously published protocols^44^. The peak height for unmodified protein and labelled protein were recorded and used to calculate percentage crosslinking using the equation:

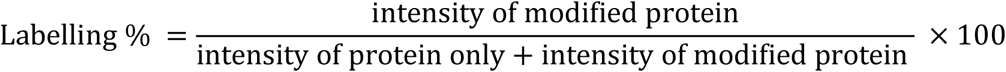

The labelling percentages at each time-point and concentration of PP12 were then plotted in GraphPad Prism 9 and fitted with one-phase association (constraints Y0 = 0). The second-order rate constant (k_inact_/K_I_) was then calculated as the slope of the linear regression fit.

### Labelling, crystallization and X-ray data collection

TYMS_26-313_ was incubated with PP12 overnight at 4 °C using a TYMS concentration of 20 μM and a TYMS_26-313_:PP12 ratio of 1:5 in 50 mM Hepes 7.5, 50 mM NaCl and 1 mM TCEP. The sample was extensively dialysed against 50 mM Hepes 7.5, 50 mM NaCl and 1 mM TCEP to remove any unbound PP12 and then concentrated to 13.5 mg/mL (400 μM) before setting up the crystallization trials. The extent of labelling, as checked by intact ESI-MS, was ≥ 92%.

Crystals of PP12-labelled TYMS26-313 were grown at 20 °C from sitting drops of 0.2 µl formed by 0.1 µl of inhibited enzyme stock solution (13.5 mg ml−1 in 50 mM Hepes 7.5, 50 mM NaCl and 1 mM TCEP) and 0.1 µl of crystallization solution containing: 10 %v/v EG, 0.1 M Hepes 7.5 pH (Buffer), 10 %v/v PEG 8K (condition 95 of Nextal PEGs II Suite, Molecular Dimensions).

Cryoprotection was provided by preparing a solution of the crystallization condition with 30% EG. The crystals were transferred into this cryoprotectant before freezing in liquid nitrogen for data collection. Diffraction data were collected at beamline I24 at the Diamond Light Source. Integration and scaling were performed using DIALS within the XIA2 expert system for X-ray diffraction data processing^72^. Crystals belong to the tetragonal space group I4_1_22 with one molecule in the crystallographic asymmetric unit.

### Structure determination and refinement

Molecular replacement was carried out with PHASER^73^, using the coordinates of human TYMS (Protain Data Bank code 6QXH)^40^ with ligands and solvent molecules removed. Models were improved and refined using COOT^74^ and PHENIX^75^. A summary of the data collection and refinement statistics is given in Supplementary Table 1. The Ramachandran plot showed 95.4 and 4.6% of residues in the most favoured and in allowed regions, respectively. Figures were drawn using PyMol (The PyMol Molecular Graphics System, v.2.0 Schrödinger, LLC).

### dTMP metabolites monitoring

#### Metabolite extraction

eHAP WT and eHAP ATRX^KO^ (g3) cells were grown to confluency in 150 mm cell culture dishes (ThermoFisher) and treated with PP12 (33 µM) or DMSO (control) for 24 hours, at which point cells were scraped and transferred to a falcon tube. Cells were washed twice with PBS and quenched on ice. Metabolite extraction was carried out by adding 500 µL of ice-cold acetonitrile/water (1/1, v/v) containing 3 nmol of ^13^C_10_,^15^N_2_ – TMP, sonicated at 4 °C (3x 8 minute pulses) and centrifuged. Supernatant was transferred to vials with glass inserts and analysed by LC-MS/MS.

#### Targeted LC-MS analysis

Samples were analysed with an adapted method as previously described^76^. Samples were injected using an Agilent Biol 1290 Infinity LC system equipped with the Bio ULD nano viper and an Hypercarb column (100 x 4.6 mm, 5 µm) from Thermo Fisher Scientific and coupled with an Agilent QQQ 6495D. Mobile phase A was composed of 0.5% (v/v) ammonium hydroxide in water containing 10 mM NH_4_HCO_3_, and mobile phase B consisted of 0.5% (v/v) ammonium hydroxide in acetonitrile/water (95/5, v/v) and 10 mM NH_4_HCO_3_. The separation of target compounds was achieved using the following gradient program at a flow rate of 0.6 mL/min: the eluting gradient started with 5% B, followed by a linear gradient to 15% B in 5.0 min, then linearly increased to 30% B in 3.0 min, to 55% in 2.0 min, returned to the initial conditions in 1.0 min to equilibrate for 4.0 min between sample injections.

MS parameters were as follows: nozzle voltage 1400 V for negative mode and capillary voltage 2.6 kV; temperature of gas and sheath gas was 225°C and 350°C; gas flow and sheath gas flow were 14.0 and 9.5 L/min, respectively. Nebulizer was set to 25 psi. Analysis was performed in dynamic multiple reaction monitoring (dMRM) mode, using the mass transitions table presented in the table below. Collision energies were optimized using commercial standards following manufacturer’s guided workflows. Samples were run in a randomised fashion, alongside pooled quality control samples and calibration curves of defined chemical standards. Qualitative and quantitative analysis were performed using Mass Hunter Qualitative Analysis and Mass Hunter Quantitative analysis (for QQQ) software (Agilent Technologies) according to the manufacturer’s workflows.

| Compound name | Precursor m/z | Product m/z | RT (min) | RT window (min) | CE (V) | Average dwell (ms) |
| --- | --- | --- | --- | --- | --- | --- |
| dTDP | 401 | 274.9 | 9.07 | 4 | 19 | 65.31 |
| dTDP | 401 | 158.9 | 9.07 | 4 | 26 | 65.31 |
| dTMP | 321.1 | 195 | 9.43 | 4 | 15 | 65.94 |
| dTMP | 321.1 | 125 | 9.43 | 4 | 22 | 65.94 |
| <sup>13</sup>C<sub>10</sub>,<sup>15</sup>N<sub>2</sub> - TMP | 377.1 | 220.8 | 9.43 | 4 | 15 | 65.94 |
| <sup>13</sup>C<sub>10</sub>,<sup>15</sup>N<sub>2</sub> - TMP | 377.1 | 185.9 | 9.43 | 4 | 15 | 65.94 |
| dTTP | 481 | 383 | 8.87 | 3 | 22 | 55.8 |
| dTTP | 481 | 158.9 | 8.87 | 3 | 26 | 55.8 |
| dUMP | 307 | 195 | 6.65 | 4 | 15 | 175.23 |
| dUMP | 307 | 111 | 6.65 | 4 | 23 | 175.23 |

### Generation of Dox-inducible Cas9 knockout cell lines

Inducible CRISPR knockout cell lines were generated by transducing eHAP iCAS9 WT and ATRX^KO^ cell lines with lentivirus produced from the lenti-sgRNA-puro construct (Addgene #104990) with the target sequence: ACCAAACGTGTGTTCTGGAA (Brunello Library – targeting Exon2 of TYMS).

To produce lentivirus, HEK293FT cells were transfected with packaging plasmids (2 µg of pLP1, 2 µg of pLP2, 2 µg of VSVG) along with 2 µg of lentiviral vector plasmid using 18 µL TransIT-Lenti (Mirus Bio) and 600 uL of Opti-MEM (Thermo Fisher Scientific).

Medium was refreshed 18 hours later and replaced with IMDM containing 10 mM Sodium Butyrate and again after an additional 6 hours with IMDM. Virus-containing supernatant was collected 48 hours post transfection, cleared through a 0.45-µm filter, supplemented with 8 µg/ml polybrene (Sigma), and used for infection of target cells. Cells were then subjected to 0.4 µg/µL puromycin selection until lentivirus-free control was no longer present (6 days). Cell lines were used as pool. Knockout efficiency was measured via western blot analysis.

### IncuCyte proliferation imaging

1000 eHAP iCas9 cells per well were seeded in opaque 96 well plates (in biological triplicates) and grown in an Incucyte S3 System (Sartorius), providing continuous live cell imaging every 4 hours up to 4 days post-seeding with a 4X objective on a full well. The “Basic Analyzer” confluence processing analysis tool was used to tailor phase segmentation and quantification.

### SDS-PAGE and immunoblotting

Proteins were separated by SDS-PAGE using NuPAGE mini gels (Invitrogen) and transferred onto 0.2 µm pore Nitrocellulose membrane (Amersham Protran; Sigma) using standard procedures. Membranes were blocked with 5% skim milk/TBST (TBS/0.1%Tween-20) for 1 hour at room temperature (RT) and probed with the indicated primary antibodies (see table below) overnight at 4°C. Membranes were then washed 3 times for 10 min with TBST, incubated with appropriate secondary antibodies conjugated to a horseradish peroxidase (HRP) for 1 hour at RT and washed again 3 times for 10 min with TBST. Immunoblots were developed using Clarity or Clarity Max Western ECL Substrate (Bio-Rad). Chemiluminescence was acquired using a ChemiDoc MP imaging system (Bio-Rad).

| Target | Dilution | Provider |
| --- | --- | --- |
| ATRX (Rabbit Polyclonal) | 1:1000 | SantaCruz (SC15408) |
| TYMS (Rabbit Monoclonal) | 1:1000 | AbCam (ab108995) |
| αtubulin (Mouse Monoclonal) | 1:10000 | Sigma (T6199) |
| H2AX (S139) (Mouse Monoclonal) | 1:1000 | Merk Millipore (05-636) |
| RPA (S33) (Rabbit Polyclonal) | 1:5000 (in milk) | Bethyl (A300264A) |

### Nuclei Count via Fluorescence Imaging

1000 eHAP iCas9 cells per well were seeded opaque 96 well plates (in biological triplicates) and grown in an Incucyte S3 System (Sartorius) for 4 days. Upon the final timepoint, plates were fixed with 4% formaldehyde for 20 minutes at RT. Plates were washed 3 times with PBS and 100 uL/well of DAPI/Triton (2 µg/µL DAPI, 0.02% Triton) solution were added and left to incubate at RT for 1h. Plates were washed 3 times with PBS and imaged on the Operetta High-Resolution Imaging microscope with a 10X objective.

### Nucleosides supplementation

Viability rescue of TYMS^KO^ cell lines has previously been reported. Protocol has been adapted from reference^77^. eHAP cells were kept in culture in 1X Iscove’s modified Dulbecco’s medium IMDM (Gibco, 21980-032) 10% Tet-Free FBS (South America origin, Fetal bovine serum, 0.2 µm sterile filtered, 500ml PAN Biotech, P30-3602) and 1% PenStrep (Gibco, 15140-122) supplemented with either dTMP (Sigma, 33430-62-5), dTTP (ThermoFisher, R0171), dT (Sigma, T1895-1G).

### Caspase III signal via Fluorescence Imaging

1500 eHAP iCas9 cells per well were seeded in opaque 96 well plates (in biological triplicates), treated as described in the text in presence of NucView 488 Caspase-3 Substrate (1:200 from 1mM stocks). Cells were grown in the Incucyte S3 System (Sartorius), providing continuous live cell imaging every 4 hours up to 4 days post-seeding with a 4X objective on a full well and recording on the Green Fluorescence Channel. The “Basic Analyzer” confluence processing analysis tool was used to tailor phase segmentation and quantification.

### Statistical analysis

Sample numbers (N or T) indicate respectively the number of independent biological samples and number of technical replicates in each experiment. Both are indicated in figure legends.

Prism 9/10 (GraphPad) and Python were used for statistical analysis: unpaired T-test was used unless stated otherwise (P < 0.05 (*), P < 0.01 (**), P < 0.001 (***), P < 0.0001 (****)). Exact P-values can be found in the figure captions.

### Additional Materials

Fragments used throughout the screen and validation were purchased from Enamine: Chemical Supplier with Catalogue Numbers: PP12: Z3766271587, PP57: Z3766270656, PP59: Z1562144202, PP2206: Z2002296328, PP1046: Z7740809535. Stereoprobes PP734 and PP735 have previously been purified as part of the work performed by *McCarthy et al*^61^.

## DATA AVAILABILITY

Crystallographic atomic coordinates and structure factors have been deposited in the PDB under accession code 31KF. The data that support the findings of this study are available from the corresponding authors upon reasonable request. Proteomics data has been deposited on the PRIDE database with the accession code PXD079268.

## ACKNOWLEDGMENTS

We would like to thank members of the Screening and Automated Science STP, Cell Science STP and Metabolomics STP at the Francis Crick Institute. We thank Andrew G. Purkiss and Simone Kunzelman of the Structural Biology Science Technology Platform at the Francis Crick Institute for assistance with diffraction data acquisition at Diamond Light Source Synchrotron (Oxford, UK) and for assistance with spectrophotometric inhibition kinetics data acquisition and analysis, respectively. The authors acknowledge I24 beamline of the Diamond Light Source Synchrotron (Oxford, UK, mx39391-34). We thank the members of the Boulton lab for discussions and valuable comments on the manuscript.

This project has been funded by a Prosperity Partnership grant from the Engineering and Physical Sciences Research Council (EPSRC), EP/ V038028/1 to [SJB, JTB, DH, AJP, MS, KR, MH]. We also thank GSK for its commitment to support fundamental discovery research through the initial establishment of the Crick-GSK LinkLabs partnership and its contribution to the EPSRC Prosperity Partnership grant.

This work was supported by the Francis Crick Institute, which receives its core funding from Cancer Research UK (CC2075, CC2000), the UK Medical Research Council (CC2075, CC2000, and the Wellcome Trust (CC2075, CC2000).

Additionally, S.S-B. was supported by an EMBO Long Term Fellowship (ALTF 707-2019) and by the European Union’s Horizon 2020 research and innovation programme under the Marie Sklodowska-Curie grant agreement No 886577.

## AUTHORS CONTRIBUTIONS

E.C., S.S.-B., S.J.B., J.T.B. and A.J.P. conceived the project, defined the ATRX-deficient synthetic-lethal phenotype, and performed the initial iteration of the phenotypic screen. M.J. performed the full phenotypic screening campaign. F.R. led the project thereafter and performed validation experiments and downstream biological studies. Chemoproteomics experiments and analyses were performed by F.R. in collaboration with E.F. Metabolomics experiments and analyses were performed by F.R. in collaboration with A.I.I. and F.T.S. C.d.C. performed crystallography and contributed to the design of *in vitro* target-validation assays. B.M.-S. and W.M. contributed to compound design and provided scientific troubleshooting and input. J.T.B. and S.J.B. supervised the project, with support from D.H., and contributed to manuscript drafting. K.R., M.H., and J.M. provided supervision and scientific input. All authors reviewed and edited the manuscript.

## DECLARATION OF CONFLICTS OF INTEREST

Federica Raguseo, Elliot Fellows, Benjamin Mortishire-Smith, David House, Andrew Powell and Jacob Bush are current or previous employees of GSK and, in most cases, GSK shareholders. There are no conflicts of interest with the work described in this publication.

## Notes

### Competing Interest Statement

The authors have declared no competing interest.

