## Supplementary Figures for "Phenotype-first covalent fragment screening identifies a synthetic lethal TYMS inhibitor in ATRX-deficient cells"

### SUPPLEMENTARY INFORMATION

| TYMS_26-313_ (PP12-inhibited)^1^ | |
| --- | --- |
| **Data collection** |  |
| Space group | I4122 |
| Cell dimensions |  |
| *a*, *b*, *c* (Å) | 110.01, 110.01, 105.09 |
| α, β, γ (°) | 90, 90, 90 |
| Resolution (Å) | 37.99 – 2.16 (2.33 – 2.16) * |
| *R*merge | 0.209 (6.744) |
| *I* / σ*I* | 12.14 (0.47) |
| Completeness (%) | 98.06 (91.09) |
| Redundancy | 26.9 (27.7) |
| **Refinement** |  |
| Resolution (Å) | 37.99-2.40 (2.486-2.400) |
| No. reflections | 17275 (3149) |
| *R*work / *R*free | 0.2442/0.2823 |
| No. atoms | 2278 |
| Protein | 2246 |
| Ligand/ion | 24 |
| Water | 8 |
| *B*-factors |  |
| Protein | 71.59 |
| Ligand/ion | 96.07 |
| Water | 64.72 |
| R.m.s. deviations |  |
| Bond lengths (Å) | 0.003 |
| Bond angles (°) | 0.62 |

*Values in parentheses are for highest-resolution shell.

^1^One crystal.

**Table 1.** Crystal Structure. Data collection and refinement statistics (molecular replacement).

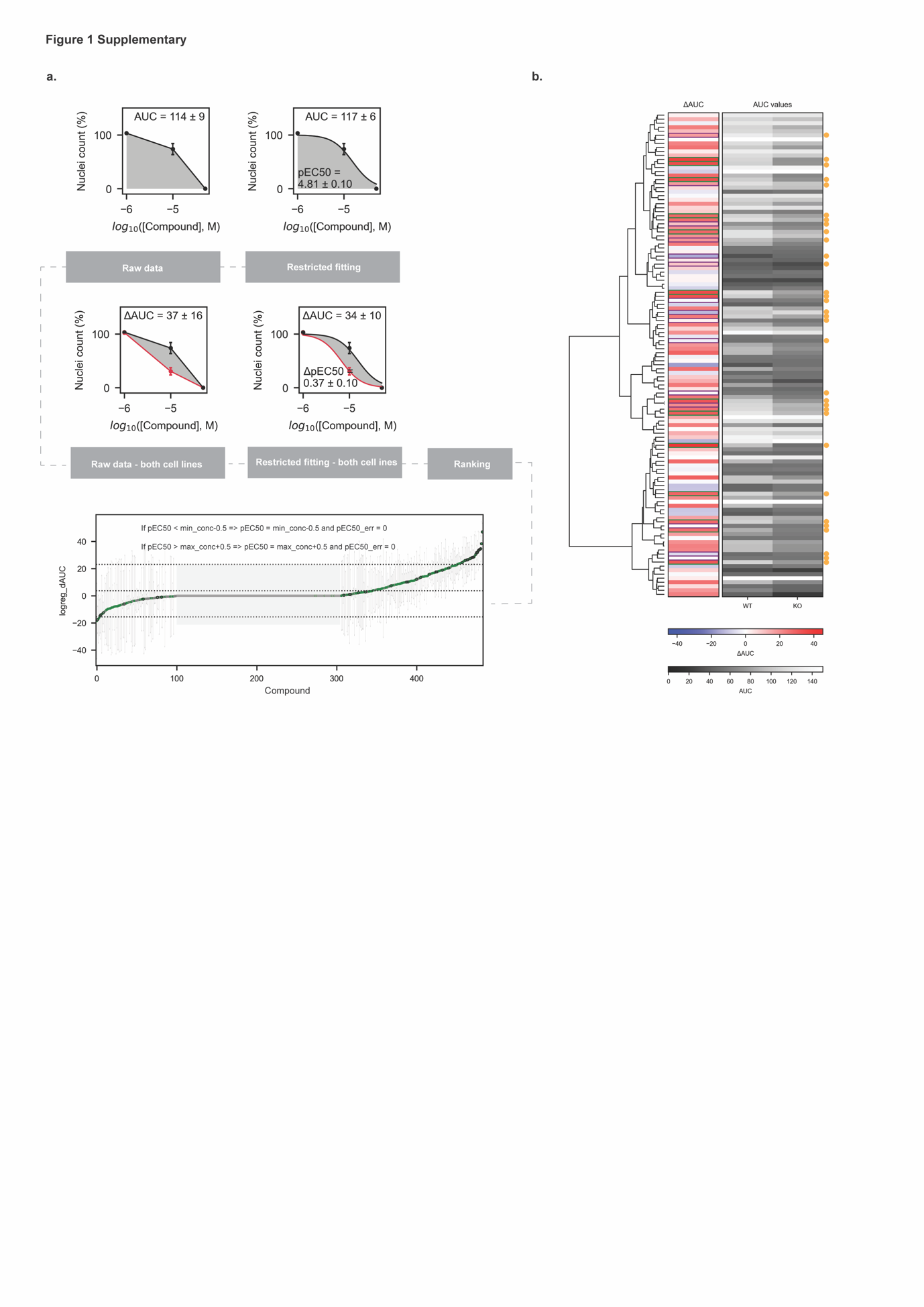

**Figure S1. High Throughput screen hit fragment deconvolution. A**. Representative hit selection analysis for compound PP63 and full ranking. Raw data is acquired as a 3-point dose response nucleic count for each cell line per compound. The data is then subjected to a restricted fitting where the highest concentration is fixed at 100% and the lowest concentration at 0% and AUC and pEC50 are extrapolated. The difference in AUC between WT and ATRX^KO^ (ΔAUC) is then calculated accordingly. All fragments are ranked according to the ΔAUC. Fragments which did not pass the filtering criteria highlighted in the figure were assigned a value of 0. **B**. Dendrogram showing structural similarities between hits. Compounds were chosen for follow-up amongst the top-ranking fragments which also presented structural similarities by using a threshold Tanimoto similarity score of ≥ 0.4. Compounds selected have been highlighted with an orange dot.

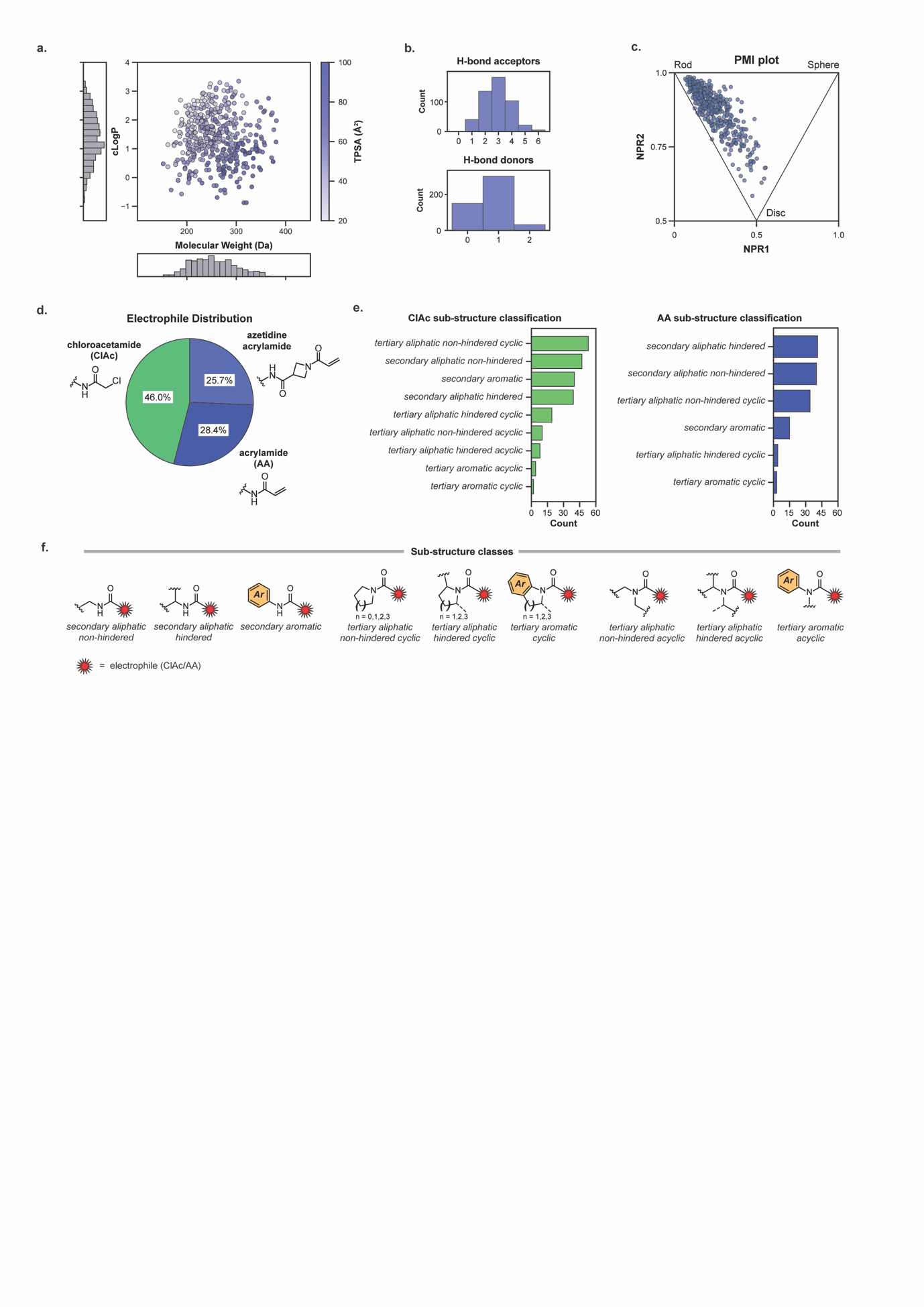

**Fig. S2. Reactive fragment library physiochemical profile**. **A**. The molecular weight and lipophilicity distributions of screened compounds. Compound lipophilicity is computed as Wildman-Crippen LogP. **B**. Distribution of H-bond acceptor and donor counts. **C**. Diversity of physical shapes in the library, based on the normalised principal moment of inertia ratios (NPR), calculated for all compounds with RDKit. **D**. Pie chart showing the electrophile distribution across the library. **E.** Sub-structure distribution of chloroacetamide (LHS) and acrylamide (RHS) functionalised fragments. Sub-structure classifications used are shown in F. **F**. SMARTS patterns classify compounds into nine sub-structures based on substitution pattern adjacent to the electrophile.

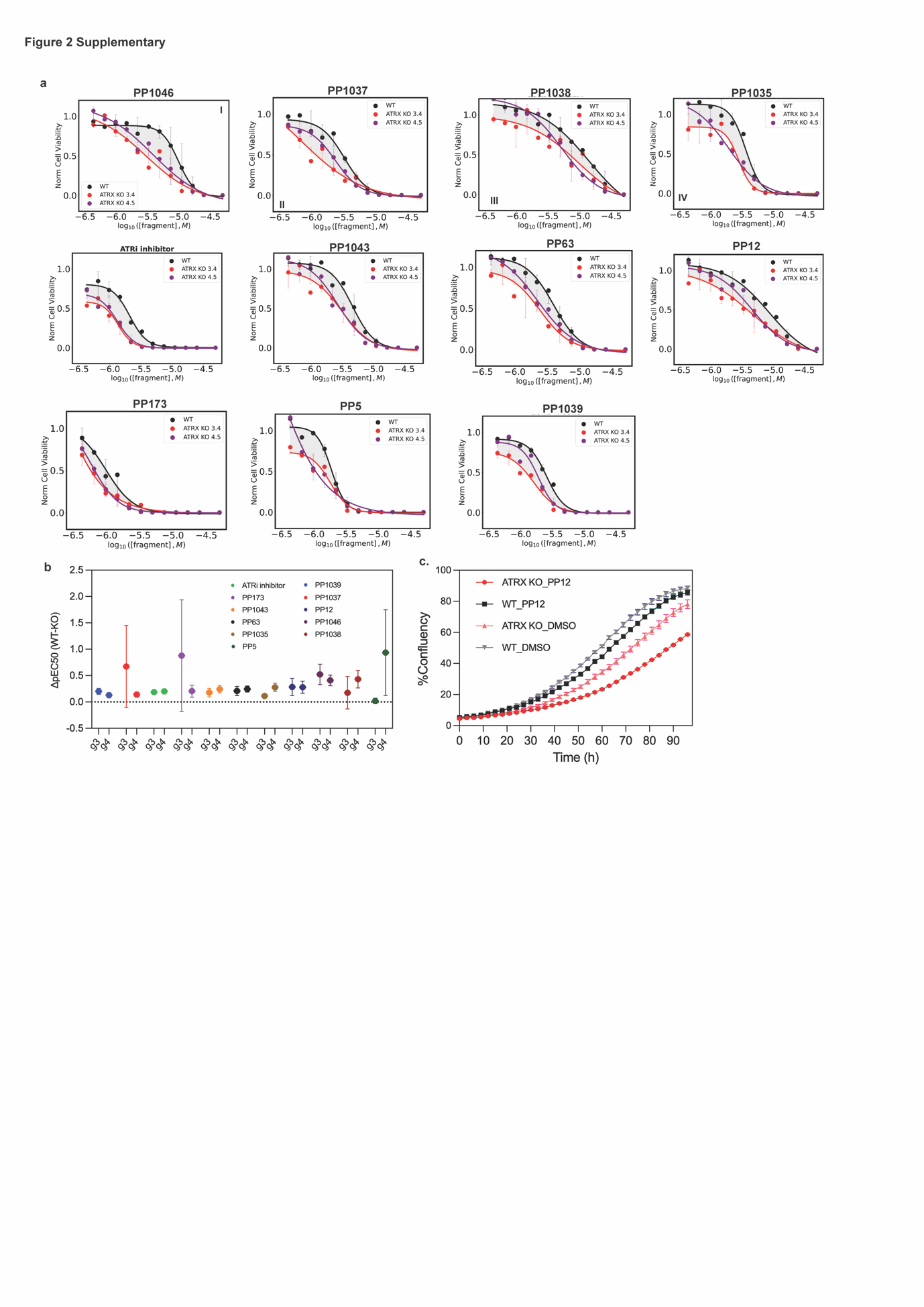

**Fig. S3. Dose-Response validation of hit fragments in presence of an additional ATRX^KO^ clone. A.** Dose-Response curves for hit fragments and alkyne-modified analogues. Cell viability was measured via CTG upon treatment of fragments in a dose-response manner. Two ATRX^KO^ clones (g3 and g4) were assessed to ensure that any observed effects were not clone-dependent. SL was compared to the ATRi positive control. N=3. **B.** Hit fragments ΔpEC50. SL phenotype was quantified for each fragment as the ΔpEC50_(WT-ATRXKO)_ for both ATRX^KO^ clones g3 and g4_._ **C.** Viability of eHAP iCAS9 WT and ATRX^KO^ upon treatment with PP12 at 6uM. Viability was further assessed in the range of the EC50 concentration (6 uM). As predicted, while the WT cell line is marginally affected with respect to the DMSO control, ATRX^KO^ growth is significantly impaired.

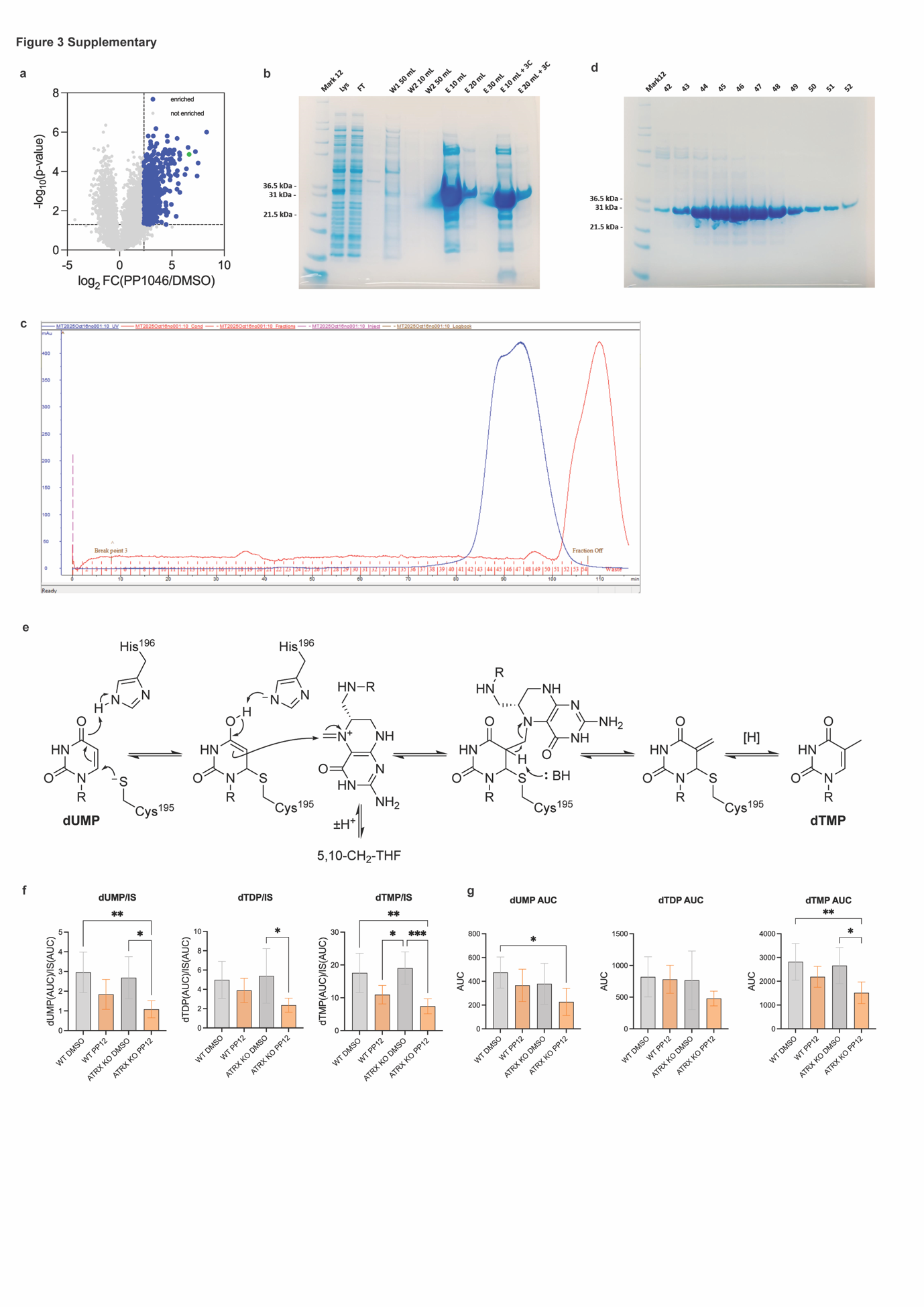

**Fig. S4. PP12 in vitro binding additional validation. A**. Protein enrichment upon PP1046 treatment. The volcano plot shows all proteins identified from the enrichment upon treatment with 50 µM PP1046 with respect to DMSO. Statistically relevant hits with over 50% enrichment are highlighted in the top right quadrant. The dataset has been overlapped with hits from the SL genetic screen performed by Segura-Bayona et al (in blue). N=3. **B.** SDS-PAGE gel of elutions upon Ni-NTA purification. TYMS protein purification elutions are reported in the gel. A band for TYMS at around 31 KDa is reported in both elutions. W=wash number. E=elution number. **C**. SEC chromatograph for TYMS purification. The peak at fractions 42-52 are representative of the TYMS protein. **D.** SDS-PAGE gel after protein purification. The TYMS band at 31 KDa is present in all fractions. Fractions with a higher concentration of TYMS (fractions from 44 to 52) were collected and concentrated for TYMS protein stocks. **E.** TYMS mechanism of action with intermediates. **F.** Targeted LC-MS analysis of dTMP, dUMP and dTDP metabolites relative to Input Signal (IS) upon treatment. **G.** Targeted LC-MS analysis of dTMP, dUMP and dTDP metabolites upon treatment reported as Area Under the Curve integration (not normalized).

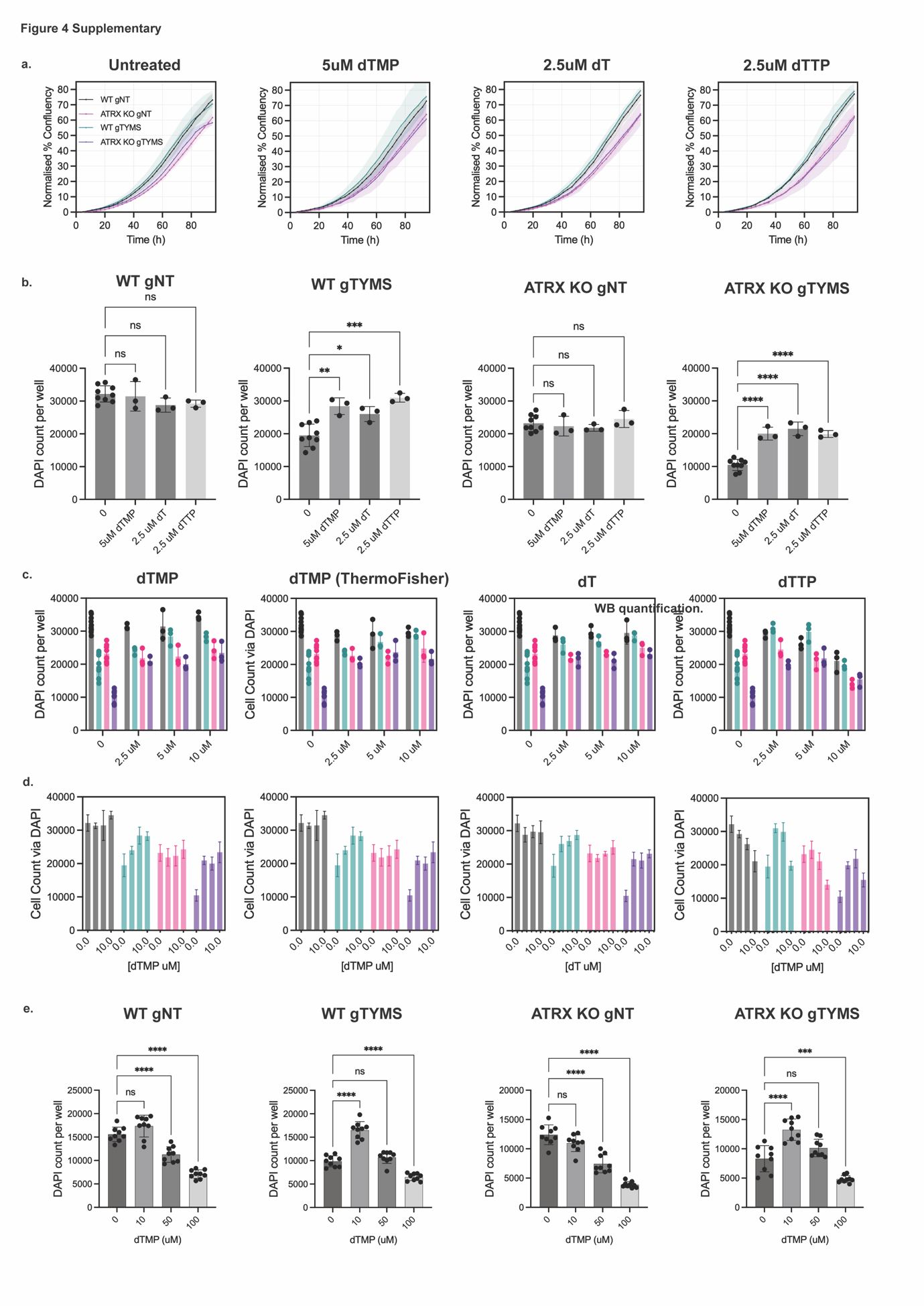

**Fig. S5. ATRX TYMS Synthetic Lethality Rescue Additional Viability Data. A**. Time-dependent growth curves for iCas9 inducible cell lines in presence of nucleosides supplementation. Viability of eHAP iCas9 ATRX^KO^ TYMS^KO^ (gTYMS), eHAP iCas9 WT TYMS^KO^ (gTYMS), eHAP iCas9 ATRX^KO^ (gNT), eHAP iCas9 WT (gNT) was measured every 4 hours via confluency measurement in the incucyte system in presence or absence of nucleosides. In untreated conditions, viability of the gTYMS cell lines is significantly impaired with respect to the gNT control, while upon supplementation the cell lines’ growth is similar with respect to the gNT control. Data was normalized with respect to confluency at t=0 for each cell line in each condition. N=3. **B**. Viability of iCas9 inducible cell lines in presence of nucleoside supplementation via DAPI count. Viability of eHAP iCas9 ATRX^KO^ TYMS^KO^ (gTYMS), eHAP iCas9 WT TYMS^KO^ (gTYMS), eHAP iCas9 ATRX^KO^ (gNT), eHAP iCas9 WT (gNT) was measured via DAPI count in presence or absence of nucleosides on day 4 after induction. Viability of the gNT cell lines is not impacted by the selected nucleosides conditions. Viability of the gTYMS cell lines increases in presence of the selected nucleosides conditions due to rescue of the phenotype. P-values are reported in the supplementary material. N=3. **C/D**. Viability of iCas9 inducible cell lines in presence of nucleoside supplementation (full curve) via DAPI count. Viability of eHAP iCas9 ATRX^KO^ TYMS^KO^ (gTYMS), eHAP iCas9 WT TYMS^KO^ (gTYMS), eHAP iCas9 ATRX^KO^ (gNT), eHAP iCas9 WT (gNT) was measured via DAPI count in presence or absence of nucleosides on day 4 after induction. Viability of the gNT cell lines is not impacted by the 0-10 µM concentration range for dTMP and dT but present a decrease in viability upon dosing with dTMP > 2.5 uM. Viability of the gTYMS cell lines increases in presence of the selected nucleosides conditions with the exception of dTMP >2.5 µM due to general toxicity effects. An additional dTMP provider (ThermoFisher) was also tested to make ensure reproducibility of the results. N=3. **E**. Viability of iCas9 inducible cell lines in presence of nucleoside supplementation (higher concentrations) via DAPI count. Viability of eHAP iCas9 ATRX^KO^ TYMS^KO^ (gTYMS), eHAP iCas9 WT TYMS^KO^ (gTYMS), eHAP iCas9 ATRX^KO^ (gNT), eHAP iCas9 WT (gNT) was measured via DAPI count in presence or absence of nucleosides on day 4 after induction. Viability of the gNT cell lines is not impacted at the 10 µM concentration of dTMP but present a decrease in viability above that threshold. Similarly, for the gTYMS cell lines, any rescue behavior is masked by toxicity above 10 µM dTMP. P-values are reported in the supplementary material. N=3.

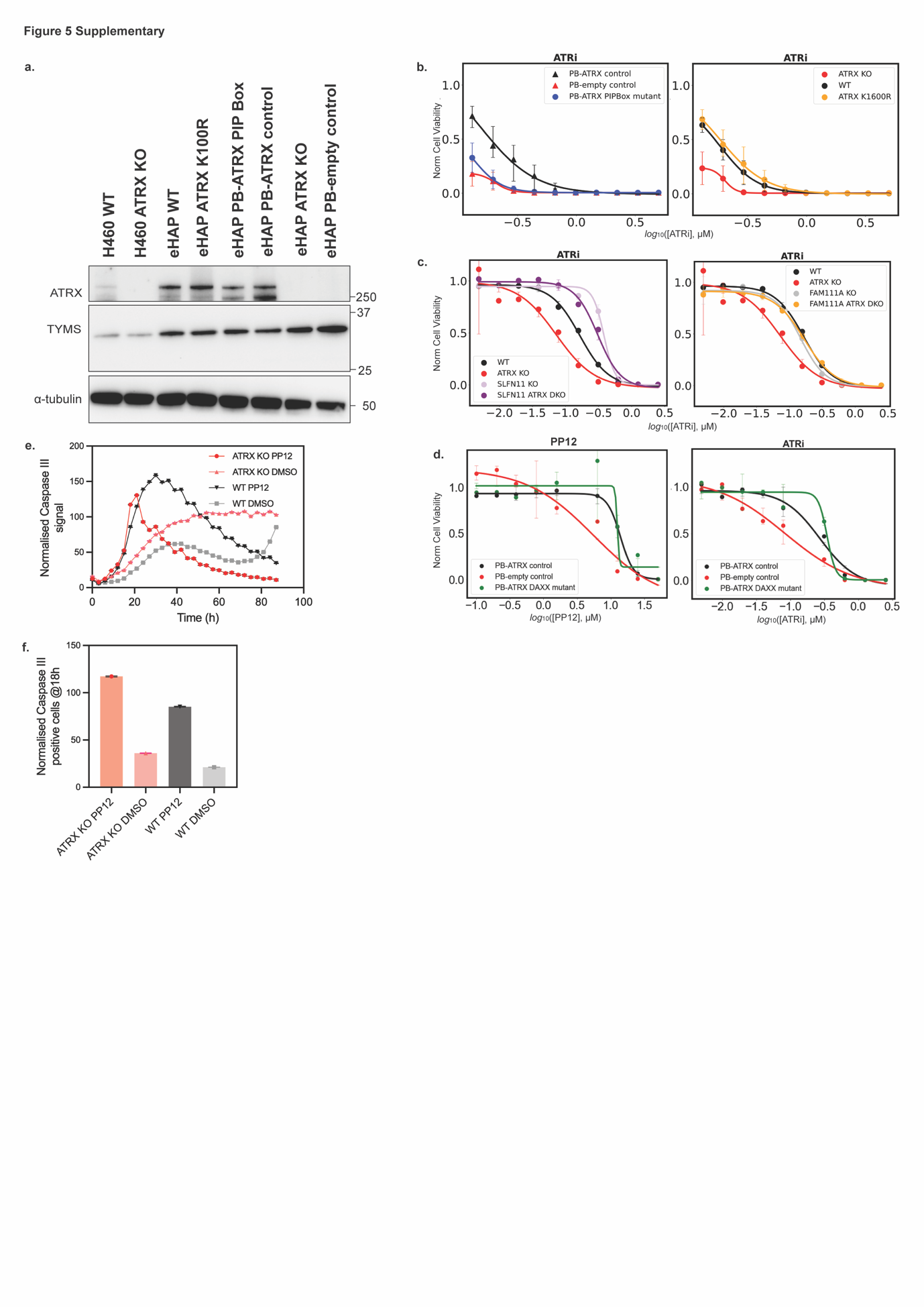

**Fig. S6. ATRX mutants experiments validation data. A**. ATRX and TYMS protein levels across mutants. Western Blot analysis was used to monitor protein levels across mutants and additional cell lines. NCI-H460 iCAS9 WT and ATRX^KO^ present comparable levels of TYMS upon ATRX-deficiency. eHAP iCAS9 ATRX^KO^ and ATRX^K100R^ present comparable levels of ATRX upon knock in. eHAP PB-ATRX^PIP box^ and PB-ATRX^control^ present comparable levels of ATRX, which are marginally lower than eHAP WT (as previously reported)^24^. **B**. Viability of mutant cell lines and controls upon treatment with ATRi positive control. Cell viability was measured via CTG upon treatment for 5 days of ATRi in a dose-response manner. The SL phenotype is still only observed for the PIP-Box mutated cell line – as in the case of PP12. N=3. **C**. Viability of double knock-out cell lines and controls upon treatment with ATRi positive control. Cell viability was measured via CTG upon treatment for 5 days of ATRi in a dose-response manner. SL is rescued upon both SLFN11 and FAM111A KO in ATRX-deficient background – as in the case of PP12. N=3. **D**. Viability of DAXX mutant cell lines and controls upon treatment with PP12 or ATRi positive control. Cell viability was measured via CTG upon treatment for 5 days of ATRi and PP12 in a dose-response manner. The SL phenotype is still not observed for the DAXX-binding domain mutated cell line for either PP12 and ATRi. N=3. **E**. Apoptosis Marker Caspase III upon acute PP12 treatment. Cells were treated with 50 µM PP12 for ~4 days and Caspase III signal was monitored in the Incucyte together with a confluency readout. Caspase III signal normalized by confluency upon time course. Both cell lines present a significant increase in Caspase III signal upon treatment. ATRX^KO^ cells are more sensitive at the 18h timepoint which is then used for follow-up experiments upon acute treatment. N=3, T=2. **F**. Apoptosis Marker Caspase III upon acute PP12 treatment at the 18h timepoint. Confluency and Caspase III signal of cells treated for 18h with 50 µM PP12 or DMSO equivalent control was recorded. In chart, the Caspase III value has been normalized by the confluency. Treated ATRX^KO^ cells present a higher Caspase III activation – implying a higher degree of apoptosis-mediated death.
